# The GET pathway serves to activate Atg32-mediated mitophagy by ER targeting of the Ppg1-Far complex

**DOI:** 10.1101/2021.09.20.461067

**Authors:** Mashun Onishi, Koji Okamoto

## Abstract

Mitophagy removes defective or superfluous mitochondria via selective autophagy. In yeast, the pro-mitophagic protein Atg32 localizes to the mitochondrial surface and interacts with the scaffold protein Atg11 to promote degradation of mitochondria. Although Atg32-Atg11 interactions are thought to be stabilized by Atg32 phosphorylation, how this posttranslational modification is regulated remains obscure. Here we show that cells lacking the guided entry of tail-anchored proteins (GET) pathway exhibit reduced Atg32 phosphorylation and Atg32-Atg11 interactions, which can be rescued by additional loss of the ER-resident Ppg1-Far complex, a multi-subunit phosphatase negatively acting in mitophagy. In GET-deficient cells, Ppg1-Far is predominantly localized to mitochondria. An artificial ER anchoring of Ppg1-Far in GET-deficient cells significantly ameliorates defects in Atg32-Atg11 interactions and mitophagy. Moreover, disruption of GET and Msp1, an AAA-ATPase that extracts non-mitochondrial proteins localized to the mitochondrial surface, elicits synthetic defects in mitophagy. Collectively, we propose that the GET pathway mediates ER targeting of Ppg1-Far, thereby preventing dysregulated suppression of mitophagy activation.

## Introduction

Mitochondria-specific autophagy, named mitophagy, is one of the membrane trafficking pathways conserved from yeast to humans. In this process, mitochondria are sequestrated by flattened double-membrane structures called isolation membranes and transported to the lysosome (in mammals) or the vacuole (in yeast), a lytic compartment, for degradation (Onishi and Okamoto, 2021; Onishi et al., 2021; Palikaras et al., 2018). In the budding yeast *Saccharomyces cerevisiae*, the outer mitochondrial membrane (OMM)-anchored protein Atg32 is phosphorylated in a manner dependent on casein kinase 2 (CK2) under mitophagy-inducing conditions (Aoki et al., 2011; Kanki et al., 2013; Kanki et al., 2009; Kondo-Okamoto et al., 2012; Okamoto et al., 2009). This posttranslational modification increases the affinity of Atg32 for Atg11, a scaffold protein for assembly of core autophagy-related (Atg) proteins required for formation of autophagosomes encapsulating mitochondria (He et al., 2006; Mao et al., 2013). Conversely, Atg32 dephosphorylation is mediated by Ppg1, a PP2A-like phosphatase (Furukawa et al., 2018). Ppg1 interacts with the Far complex that acts in a cooperative manner to suppress Atg32 phosphorylation, Atg32-Atg11 interactions, and mitophagy (Furukawa et al., 2018). Together, the phosphorylation-dephosphorylation switch for Atg32 is likely to be a key regulatory step to initiate selective degradation of mitochondria.

Appropriate targeting of membrane proteins to correct subcellular destinations is critical to maintain functional compartments within cells (Barlowe and Miller, 2013). Tail-anchored (TA) proteins, which harbor a single transmembrane (TM) domain at the very C-terminus, are post-translationally inserted into the membranes of mitochondria, peroxisomes, and ER, acting in a myriad of cellular processes such as vesicular trafficking, protein import, and organelle dynamics (Barlowe and Miller, 2013). In budding yeast, multiple TA proteins are targeted to the ER via the guided entry of TA proteins (GET) pathway (Denic, 2012; Denic et al., 2013; Farkas and Bohnsack, 2021). Prior to insertion into the ER membrane, the TM domains of TA proteins are shielded by the cytosolic ATPase Get3 (Bozkurt et al., 2009; Mateja et al., 2015; Mateja et al., 2009; Suloway et al., 2009; Yamagata et al., 2010). Then, the Get3-TA protein complexes are recruited to the ER membrane-embedded Get1/2 insertase complex (McDowell et al., 2020; Schuldiner et al., 2008; Stefer et al., 2011; Wang et al., 2014; Wang et al., 2011). Successful interactions between Get1/2 and Get3 drive detachment of TA proteins from Get3, enabling their insertion into the ER membrane by the Get1/2 complex. Upon disruption of the GET pathway, several TA proteins are not properly localized to the ER, but instead, targeted to mitochondria (Jonikas et al., 2009; Schuldiner et al., 2008). These ER-resident TA proteins on the OMM are removed by Msp1, a mitochondrial surface-anchored AAA-ATPase that extracts inappropriately targeted non-mitochondrial TA proteins and thus maintains mitochondrial membrane integrity (Chen et al., 2014; Okreglak and Walter, 2014; Wang et al., 2020; Wohlever et al., 2017; Zhang et al., 2011).

Our previous findings reveal a previously unappreciated role for Get1/2 in promoting mitophagy during prolonged respiratory growth (Onishi et al., 2018). In contrast to severely impaired mitophagy, other selective and bulk autophagy pathways are only slightly affected, indicating that the common core autophagy machinery itself is rarely altered in the absence of Get1/2 (Onishi et al., 2018). Although it is likely that the Get1/2 complex serves a specialized function in mitophagy, how this ER-resident TA protein insertase acts in degradation of mitochondria remains uncertain. In this study, we demonstrate that Atg32 phosphorylation and Atg32-Atg11 interactions are compromised in cells lacking Get components. Notably, perturbation of Ppg1-mediated Atg32 dephosphorylation mostly recovers Atg32-Atg11 interactions and mitophagy in *get1/2*-null cells. Moreover, the Ppg1-Far complex is localized to the ER in a manner dependent on the GET pathway, and loss of the Get components leads to targeting of this phosphatase complex to mitochondria. Artificial ER localization of the Far complex in the absence of Get1/2 significantly restores Atg32-Atg11 interactions and mitophagy. In addition, disruption of Msp1 extractase activity in GET-deficient cells causes an exacerbation in mitophagy defects. Taken together, our data suggest that the GET pathway serves to promote appropriate targeting of the Ppg1-Far complex to the ER, thereby contributing to Atg32 activation at the initial stage of mitophagy.

## Results

### Atg32 phosphorylation and Atg32-Atg11 interactions are reduced in cells lacking Get components

In yeast, mitophagy initiation consists of three main steps, expression, mitochondrial localization, and phosphorylation of Atg32. Based on our previous results that Get components are not critical for Atg32 expression and mitochondrial localization (Onishi et al., 2018), we sought to test if loss of Get components affects Atg32 phosphorylation in the early phase of mitophagy. Atg32 is phosphorylated when wild-type cells are grown in respiratory media containing non-fermentable carbon sources such as glycerol (Gly) (Kondo-Okamoto et al., 2012). Under these mitophagy-inducing conditions, putative phosphorylated Atg32 molecules appeared as multiple upper bands (Fig. 1 A) that were diminished by treatment with a protein phosphatase (Fig. 1 B). In contrast, these mobility shifts seemed to be reduced in *get1/get2/get3*-null cells, indicating that Get components are important for efficient phosphorylation of Atg32 (Fig. 1 A).

**Figure 1.**
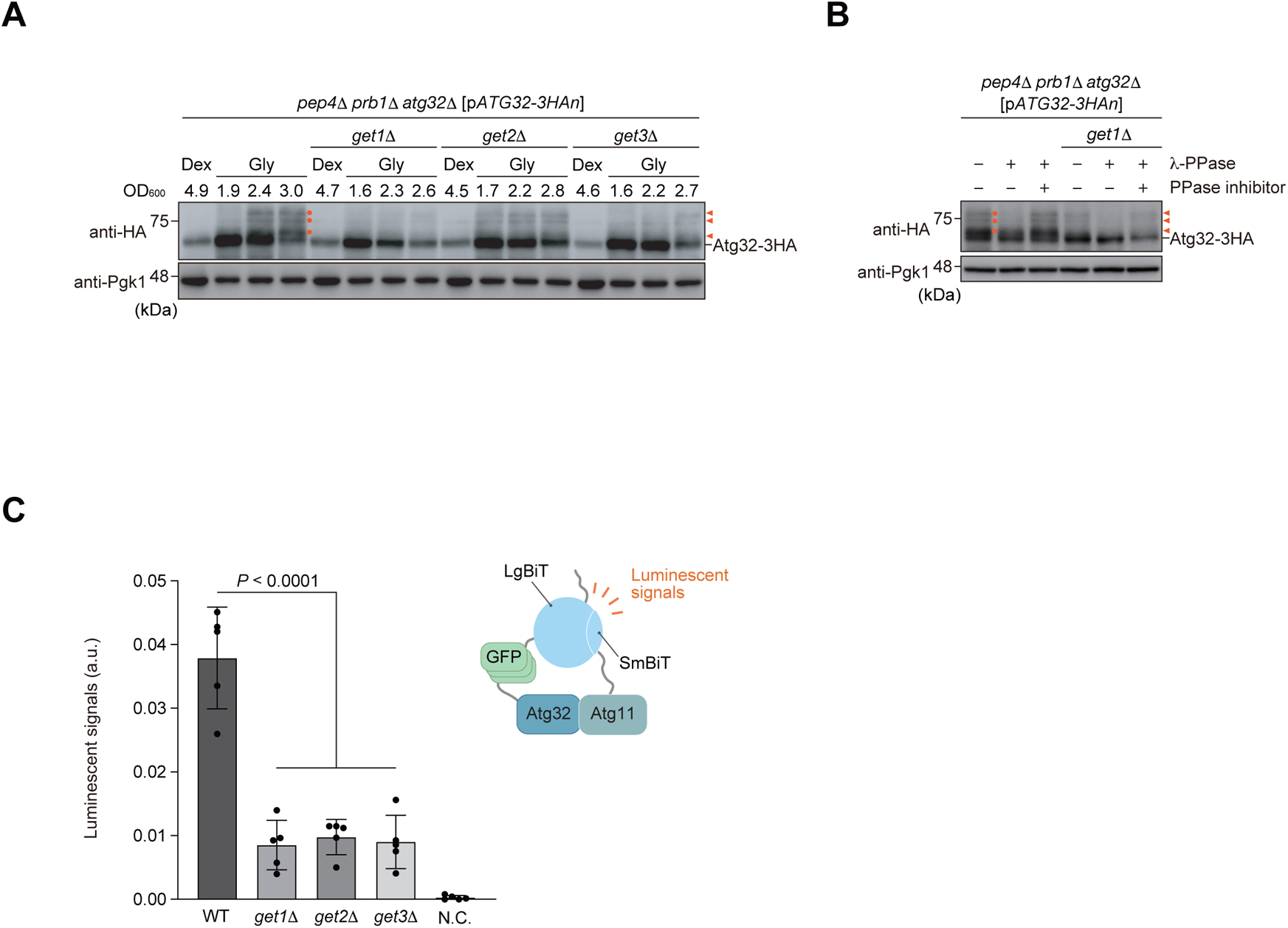
Atg32 phosphorylation and Atg32-Atg11 interactions are reduced in cells lacking Get components. **(A)** Wild-type, *get1Δ*, *get2Δ*, and *get3*Δ cells containing a plasmid encoding Atg32-3HAn (p-*ATG32-3HAn*) grown in fermentable dextrose medium (Dex) were cultured in non-fermentable glycerol medium (Gly), collected at the indicated OD600 points, and subjected to western blotting. All strains are *pep4/prb1/atg32*-triple-null derivatives defective in vacuolar degradation of Atg32-3HAn via mitophagy. Atg32 is phosphorylated at the early stages of respiratory growth, and phosphorylated Atg32 molecules are detected as multiple upper protein bands. Orange arrowheads and dots indicate putative phosphorylated Atg32. Pgk1 was monitored as a loading control. **(B)** *pep4*Δ *prb1*Δ *atg32*Δ and *pep4*Δ *prb1*Δ *atg32*Δ *get1*Δ cells containing a plasmid encoding Atg32-HAn were grown in glycerol medium, collected at the OD600 = 2.5 point, and subjected to alkaline lysis and TCA precipitation. The pellet was resuspended in a reaction buffer, treated with or without lambda protein phosphatase (λ-PPase) in the presence or absence of PPase inhibitor. **(C)** Wild-type, *get1*Δ, *get2*Δ and *get3*Δ cells expressing Atg32 internally tagged with 3×GFP plus Large BiT (LgBiT) and Atg11 C-terminally tagged with Small BiT (SmBiT) or wild-type cells expressing Atg32 and Atg11 (negative control, N.C.), were grown in glycerol medium, collected at the OD600 = 1.4 point, incubated with substrates, and subjected to the NanoBiT-based bioluminescence assay. Data represent the averages of all experiments, with bars indicating standard deviations (*n* = 5 independent cultures). Data were analyzed by one-way analysis of variance (ANOVA) with Dunnett’s multiple comparison test.

As Atg32 phosphorylation is thought to be a key regulatory step for stabilizing Atg32-Atg11 interactions (Aoki et al., 2011; Kondo-Okamoto et al., 2012), we next investigated whether loss of Get components impinges this protein-protein interaction for mitophagy. To address this issue, we applied the NanoBiT (NanoLuc Binary Technology, Promega) system, a luminescence-based assay for protein-protein interactions, to quantitative monitoring of Atg32-Atg11 interactions in live cells. When yeast cells expressing chromosomally integrated LgBiT-tagged Atg32 and SmBiT-tagged Atg11 were grown under respiratory conditions, the Atg32-Atg11 interaction brings the LgBiT and SmBiT subunits into close proximity, resulting in reversible reconstitution of an active luciferase that generates a luminescent signal in the presence of its substrate furimazine (Dixon et al., 2016) (Fig. S1 A). This system, which efficiently drives mitophagy (80% compared to wild-type cells) without overexpression, enables us to measure the resulting luminescent signals by a microplate reader and relatively quantify Atg32-Atg11 interactions *in vivo*. Our NanoBiT system detected lower luminescent signals in cells lacking Get1, Get2, or Get3 (3-5-fold reduction compared to wild-type cells) under respiratory conditions (Fig. 1 C), indicating that Get components are required for promoting Atg32-Atg11 interactions.

### Perturbation of the Ppg1 phosphatase restores Atg32-Atg11 interactions and mitophagy in get1/2-null cells

It is conceivable that a decrease in Atg32 phosphorylation causes suppression of Atg32-Atg11 interactions in cells lacking Get components (Fig. 1, A and C). Thus, we hypothesized that augmentation of Atg32 phosphorylation could rescue the impaired protein-protein interactions for mitophagy in GET-deficient cells. To test this possibility, we attempted to genetically increment Atg32 phosphorylation by loss of Ppg1, a protein phosphatase acting in dephosphorylation of Atg32 and suppression of Atg32-Atg11 interactions (Furukawa et al., 2018). Accordingly, we performed the NanoBiT assay and found that consistent with the previous report (Furukawa et al., 2018), loss of Ppg1 increased Atg32-Atg11 interactions (2-3-fold compared to wild-type cells) (Fig. 2 A). Remarkably, in *get1/2 ppg1-*double-null cells, Atg32 interacted with Atg11 at near wild-type levels, supporting the idea that reduced Atg32 phosphorylation in cells lacking Get1/2 is the primary cause of a defect in Atg32-Atg11 interactions (Fig. 2 A).

**Figure 2.**
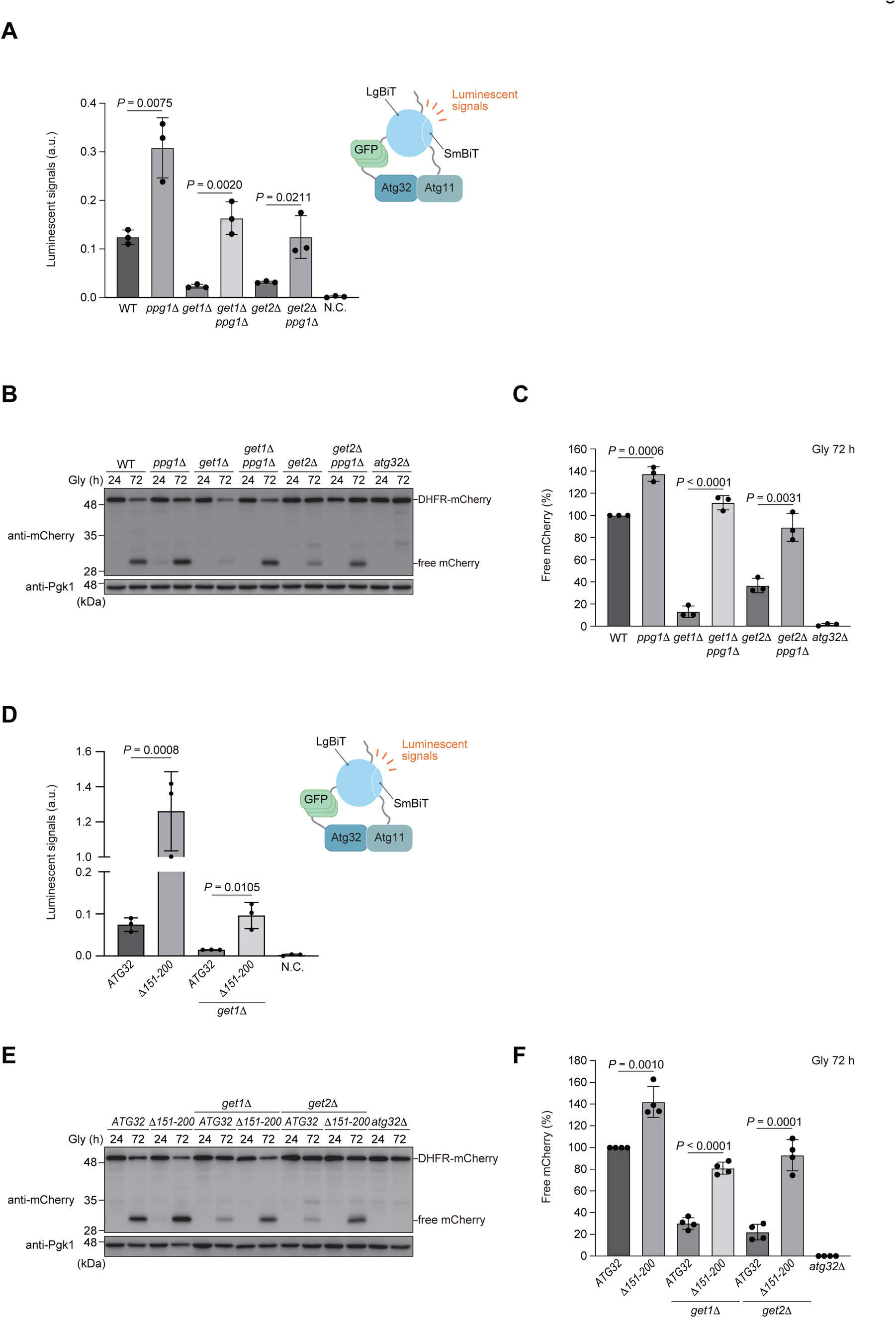
Perturbation of the Ppg1 phosphatase restores Atg32-Atg11 interactions and mitophagy in *get1/2*-null cells. **(A)** Wild-type, *ppg1*Δ, *get1*Δ, *get2*Δ, *get1*Δ *ppg1*Δ, and *get2*Δ *ppg1*Δ cells expressing Atg32-3HA-3×GFP-3FLAG-LgBiT and Atg11-HA-SmBiT, or wild-type cells expressing Atg32 and Atg11 (negative control, N.C.) were grown in glycerol medium (Gly), collected at the OD600 = 1.4 point, incubated with substrates, and subjected to the NanoBiT-based bioluminescence assay. Data represent the averages of all experiments, with bars indicating standard deviations (*n* = 3 independent cultures). **(B)** Wild-type, *ppg1*Δ, *get1*Δ, *get2*Δ, *get1*Δ *ppg1*Δ, *get2*Δ *ppg1*Δ, and *atg32*Δ cells expressing mitochondrial matrix-targeted DHFR-mCherry (mito-DHFR-mCherry) were grown in glycerol medium (Gly), collected at the indicated time points, and subjected to western blotting. Generation of free mCherry indicates transport of mitochondria to the vacuole. **(C)** The amounts of free mCherry in cells analyzed in **(B)** was quantified in three experiments. The signal intensity value of free mCherry in wild-type cells at the 72 h time point was set to 100%. Data represent the averages of all experiments, with bars indicating standard deviations (*n* = 3 independent cultures). **(D)** Wild-type and *get1*Δ cells expressing Atg11-HA-SmBiT and Atg32-3HA-3×GFP-3FLAG-LgBiT or (Δ151-200)-3HA-3×GFP-3FLAG-LgBiT, or wild-type cells expressing Atg32 and Atg11 (negative control, N.C.) were grown in glycerol medium (Gly), collected at the OD600 = 1.4 point, incubated with substrates, and subjected to the NanoBiT-based bioluminescence assay. Data represent the averages of all experiments, with bars indicating standard deviations (*n* = 3 independent cultures). **(E)** Wild-type, *get1*Δ, and *get2*Δ cells expressing chromosomally integrated *ATG32* wild-type or *ATG32 (Δ151-200)* were grown in glycerol medium (Gly), collected at the indicated time points, and subjected to western blotting. **(F)** The amounts of free mCherry in cells analyzed in **(E)** was quantified in four experiments. The signal intensity value of free mCherry in wild-type cells at the 72 h time point was set to 100%. Data represent the averages of all experiments, with bars indicating standard deviations (*n* = 4 independent cultures). Data were analyzed by two-tailed Student’s *t* test **(A, C, D, F)**.

Next, we performed mitophagy assay using a mitochondrial matrix-localized DHFR-mCherry (mito-DHFR-mCherry) probe (Calvelli et al., 2020). When mitochondria are transported to the vacuole, DHFR-mCherry is processed by vacuolar proteases to generate free mCherry, enabling semi-quantitative detection of mitochondrial degradation. We confirmed that loss of Ppg1 accelerated mitophagy (137% compared to wild-type cells) (Fig. 2, B and C). Strikingly, *get1/ppg1-* and *get2/ppg1*-double-null cells exhibited mitophagy at near wild-type levels (112% and 89%, respectively, compared to wild-type cells) (Fig. 2, B and C). Moreover, expression of a *PPG1 H111N* gene encoding a catalytically inactive phosphatase restored Atg32-Atg11 interactions and mitophagy in *get1/*2-null cells (Fig. S1, B-D). Together, these results suggest that perturbation of Ppg1 increased the affinity of Atg32 for Atg11 in *get1/*2-null cells, thereby recovering mitophagy.

To exclude the possibility that restoration of mitophagy in *get1/ppg1-* and *get2/ppg1*-double-null cells is caused indirectly by pleiotropic alterations in Ppg1 substrate(s), we examined Atg32 variants lacking the amino acid residues 151-200 that are required for the Ppg1-Far complex to interact with Atg32 (Furukawa et al., 2018; Innokentev et al., 2020). When this truncation was introduced into the NanoBiT system, the Atg32 mutant (Δ151-200) interacted with Atg11 14-18-fold more strongly than the full-length protein in the presence of Get1, and at near wild-type levels even in the absence of Get1 (Fig. 2 D). Consistent with these results, mitophagy in *get1/2*-null cells were mostly restored by expression of the Atg32 mutant (Δ151-200) (Fig. 2, E and F), supporting the idea that Ppg1-Far-mediated suppression of Atg32-Atg11 interactions and mitophagy is exacerbated in the absence of Get1/2.

### The ER-resident Far complex predominantly targets to mitochondria in GET-deficient cells

How could Ppg1 abrogate mitophagy in cells lacking Get1/2? It has been demonstrated that ER-resident TA proteins localize to mitochondria in *get1/2*-null cells (Jonikas et al., 2009; Schuldiner et al., 2008). In addition, Ppg1 interacts with the Far complex that acts in pheromone-induced cell cycle arrest and the TORC2 signaling pathway (Furukawa et al., 2018; Kemp and Sprague, 2003; Pracheil et al., 2012). Moreover, the Far complex contains the TA proteins Far9 and Far10, and is anchored to the ER membrane in a manner dependent on their TA domains (Pracheil and Liu, 2013). Based on these findings, we hypothesized that disruption of the GET pathway may lead to targeting of the ER-resident Ppg1-Far complex to the surface of mitochondria, thereby oversuppressing mitophagy. To test this idea, Far8, a component of the Far complex, was functionally tagged with three copies of GFP, expressed from the chromosomal *FAR8* locus without overexpression, and observed using fluorescence microscopy. We found that Far8-3×GFP mostly colocalized with mCherry-tagged Sec63, an ER-anchored Hsp40/DnaJ family protein (Feldheim et al., 1992) that exhibited peripheral and perinuclear patterns, in wild-type cells under respiratory conditions (Fig. 3, A and B). By contrast, Far8-3×GFP predominantly localized to mitochondria in *get1*-null cells (93% of cells lacking Get1 and 9% of wild-type cells) (Fig. 3, C and D). We also confirmed that loss of Get2 or Get3 greatly increased mitochondria-targeted Far8-3×GFP (96% and 92% of *get2*- and *get3*-null cells, respectively) (Fig. S2 A).

**Figure 3.**
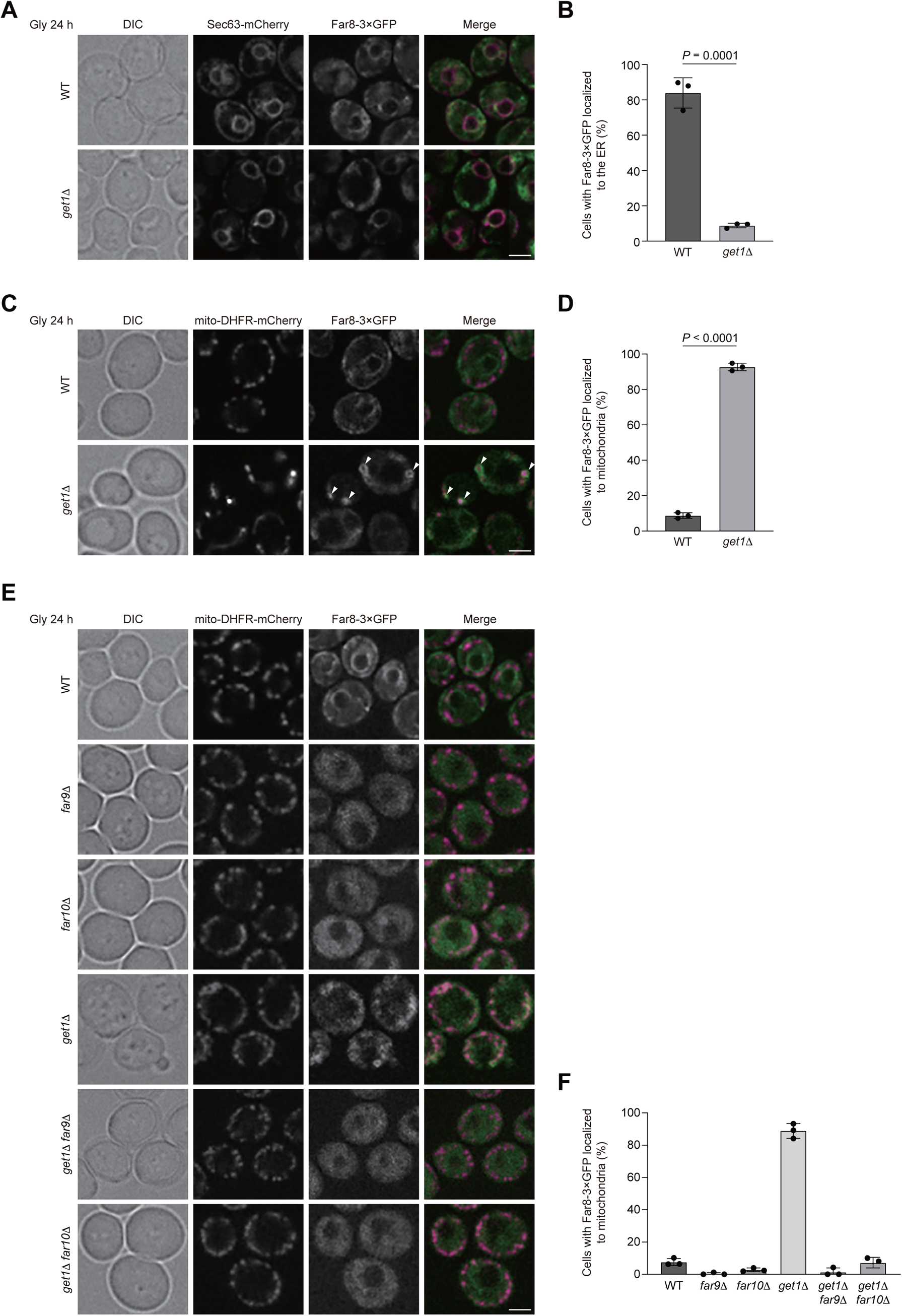
The ER-resident Far complex predominantly targets to mitochondria in GET-deficient cells. **(A)** Representative images of wild-type and *get1*Δ cells expressing Sec63-mCherry and Far8-3×GFP grown for 24 h in glycerol medium (Gly) and observed by structured illumination microscopy. Arrowheads indicate Far8-3×GFP localized to mitochondria. Scale bar, 2 μm. DIC, differential interference contrast. **(B)** Cells analyzed in **(A)** were quantified in three experiments. Data represent the averages of all experiments, with bars indicating standard deviations (*n* = 3 independent cultures). **(C)** Representative images of wild-type and *get1*Δ cells expressing mito-DHFR-mCherry and Far8-3×GFP grown for 24 h in glycerol medium (Gly) and observed by structured illumination microscopy. Scale bar, 2 μm. **(D)** Cells analyzed in **(C)** were quantified in three experiments. Data represent the averages of all experiments, with bars indicating standard deviations (*n* = 3 independent cultures). **(E)** Representative images of wild-type, *far9*Δ, *far10*Δ, *get1*Δ, *get1*Δ *far9*Δ, and *get1*Δ *far10*Δ cells expressing Sec63-mCherry and Far8-3×GFP3 grown for 24 h in glycerol medium (Gly) and observed by structured illumination microscopy. Scale bar, 2 µm. **(F)** Cells analyzed in **(E)** were quantified in three experiments. Data represent the averages of all experiments, with bars indicating standard deviations. Data were analyzed by two-tailed Student’s *t* test **(B, D)**.

To clarify whether the insertase activity of Get1/2 is required for Far8-3×GFP localization to the ER, we generated yeast strains expressing an inactive Get1 or Get2 variant with point mutations in their conserved cytosolic domain (Get1NRm: N72A, R73A, Get2RERRm: R14E, E15R, R16E, R17E) (Wang et al., 2011), and found that expression of these insertase-inactive mutants significantly disturbed ER localization of Far8-3×GFP (95% and 98% of cells expressing Get1NRm and Get2RERRm, respectively) (Fig. S2 B), further underscoring a primary role for the GET pathway in ER targeting of the Ppg1-Far complex. In cells expressing these mutants, mitophagy was moderately reduced (Fig. S2, C and D), indicating that the Get1/2 insertase activity is required for efficient mitophagy.

Since targeting of ER-resident TA proteins to the mitochondrial surface requires their TA domains (Farkas and Bohnsack, 2021), we assumed that loss of Far9 or Far10 could diminish mitochondrial localization of the Far complex in cells lacking Get1/2. In line with this idea, we found that Far8-3×GFP was hardly localized to mitochondria, but instead mostly dispersed throughout the cytoplasm (probably excluded from the vacuolar lumen) in *far9/10*-null, *far9/get1*- and *far10/get1*-double-null cells (Fig. 3, E and F), indicating that these TA proteins are indispensable for targeting of the Far complex to the ER in wild-type cells or mitochondria in GET-deficient cells.

It has recently been reported that a fraction of the Far complex is localized to mitochondria even in wild-type cells under fermentable conditions (Innokentev et al., 2020). Although we barely found mitochondrial localization of Far8-3×GFP under non-fermentable conditions (Fig. 3, A and C), it remained possible that a small fraction of the Far complex is localized to mitochondria and degraded in a mitophagy-dependent manner. To clarify this issue, we performed GFP-processing assays. Similar to mito-DHFR-mCherry, Far8-3×GFP localized to the ER and mitochondria can be transported to the vacuole and processed to generate free GFP via ER-phagy and mitophagy, respectively. Under respiratory conditions, generation of free GFP was reduced by 50% in cells without mitophagy (*atg32*-null) or ER-phagy (*atg39/40*-double-null) (Mochida et al., 2015) and 25% in cells without both events (*atg32/39/40*-triple-null) compared to wild-type cells (Fig. S2, E and F). These results support the notion that a small fraction of the Ppg1-Far complex escapes the GET pathway and localizes to the surface of mitochondria.

### Loss of the Far9/10 TA proteins rescues mitophagic deficiencies in cells lacking Get1/2

Our observations that mitochondrial localization of the Far complex in *get1*-null cells was diminished by loss of Far9 or Far10 (Fig. 3, E and F) led us to examine Atg32-Atg11 interactions and mitophagy in the absence of these TA proteins. Similar to the results obtained from *ppg1*-null cells (Fig. 2, A-C), Atg32 interacted with Atg11 2-3-fold more strongly in cells lacking Far9 than wild-type cells (Fig. 4 A). In addition, consistent with the previous findings (Furukawa et al., 2018), mitophagy under respiratory conditions was increased in *far9*-null cells (139% compared to wild-type cells) (Fig. 4, B and C). Strikingly, Atg32-Atg11 interactions and mitophagy were restored at near wild-type levels in *get1/far9*- and *get2/far9*-double-null cells (Fig. 4, A-C).

**Figure 4.**
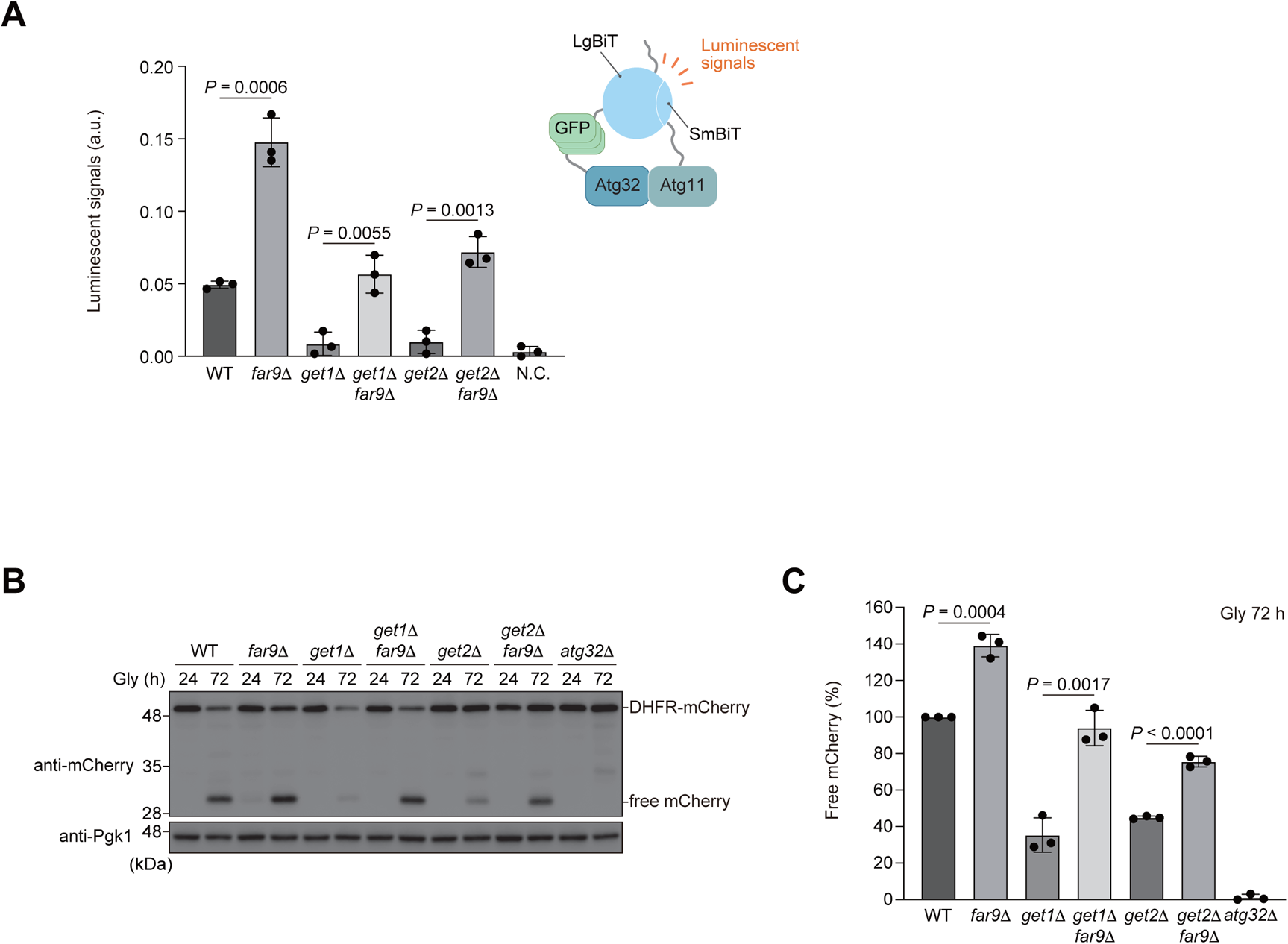
Loss of the Far9/10 TA proteins rescues mitophagic deficiencies in cells lacking Get1/2. **(A)** Wild-type, *far9*Δ, *get1*Δ, *get2*Δ, *get1*Δ *far9*Δ, and *get2*Δ *far9*Δ cells expressing Atg32-3HA-3×GFP-3FLAG-LgBiT and Atg11-HA-SmBiT, or wild-type cells expressing Atg32 and Atg11 (negative control, N.C.) were grown in glycerol medium (Gly), collected at the OD600 = 1.4 point, incubated with substrates, and subjected to the NanoBiT-based bioluminescence assay. Data represent the averages of all experiments, with bars indicating standard deviations (*n* = 3 independent cultures). **(B)** Wild-type, *far9*Δ, *get1*Δ, *get2*Δ, *get1*Δ *far9*Δ, *get2*Δ *far9*Δ, and *atg32*Δ cells expressing mito-DHFR-mCherry were grown in glycerol medium (Gly), collected at the indicated time points, and subjected to western blotting. **(C)** The amounts of free mCherry in cells analyzed in **(B)** was quantified in three experiments. The signal intensity value of free mCherry in wild-type cells at the 72 h time point was set to 100%. Data represent the averages of all experiments, with bars indicating standard deviations (*n* = 3 independent cultures). Data were analyzed by two-tailed Student’s *t* test **(A, C)**.

Next, we investigated cells lacking Far10 and found only a slight and no increase in Atg32-Atg11 interactions and mitophagy (1.2-fold and 98%, respectively, compared to wild-type cells) (Fig. S3, A-C). Notably, *get1/far10*- and *get2/far10*-double-null cells exhibited a partial recovery in Atg32-Atg11 interactions (0.7- and 0.4-fold, respectively, compared to wild-type cells) (Fig. S3 A) and a substantial restoration in mitophagy (96% and 69%, respectively, compared to wild-type cells) (Fig. S3, B and C). Collectively, these data suggest that the Ppg1-Far complex is anchored to the mitochondrial surface via Far9/10 and acts in suppression of Atg32-Atg11 interactions and mitophagy.

### Artificial ER anchoring of the Far complex increases Atg32-Atg11 interactions and mitophagy in get1/2-null cells

Based on our findings that loss of Get1/2 leads to excess mitochondrial localization of the Ppg1-Far complex (Fig. 3, A-D; and Fig. S2 A), we asked whether Get1/2-independent ER localization of the Ppg1-Far complex ameliorates mitophagy deficiencies in cells lacking Get1/2. To this end, the TA domain of Far9 was replaced with the TM domain (TM^ER^) of Sec12, a single-pass ER membrane protein consisting of an N- and C-terminal domains facing the cytosol and ER lumen, respectively (d’Enfert et al., 1991). We confirmed that expression of Far9-TM^ER^ does not cause significant alterations in ER shape and Far8-3×GFP localizations (Fig. 5, A and B). As expected, Far8-3×GFP in cells expressing Far9-TM^ER^ was localized to the ER even in *get1/2*-null cells (Fig. 5, A and B; and Fig. S4, A and B). In addition, expression of Far9-TM^ER^ in cells lacking Get1/2 restored Atg32-Atg11 interactions at near wild-type levels (Fig. 5 C). Moreover, mitophagy was increased in *get1*- and *get2*-null cells (70% and 80%, respectively compared to wild-type cells) (Fig. 5, D and E), suggesting that ER retention of the Ppg1-Far complex is critical for efficient mitophagy.

**Figure 5.**
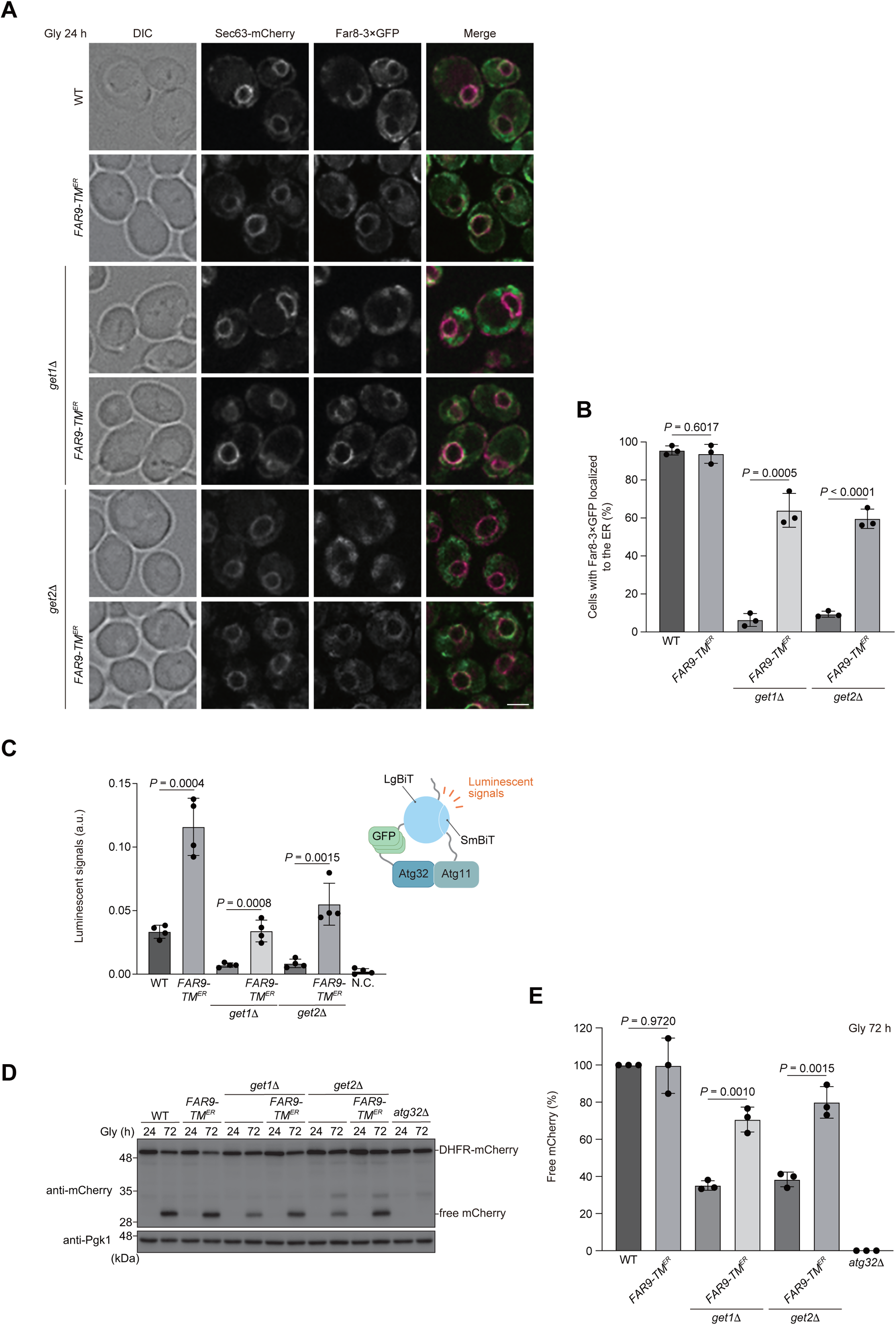
Artificial ER anchoring of the Far complex increases Atg32-Atg11 interactions and mitophagy in *get1/2*-null cells. **(A)** Representative images of wild-type, *get1*Δ, and *get2*Δ cells expressing the endogenous Far9 or a variant whose TA domain was replaced with the Sec12 TM domain (*FAR9-TM^ER^*) grown for 24 h in glycerol medium (Gly) and observed by structured illumination microscopy. All strains were derivatives expressing Sec63-mCherry and Far8-3×GFP. Scale bar, 2 μm. **(B)** Cells analyzed in **(A)** were quantified in three experiments. Data represent the averages of all experiments, with bars indicating standard deviations (*n* = 3 independent cultures). **(C)** Derivatives of cells analyzed in **(D)** expressing Atg32-3HA-3×GFP-3FLAG-LgBiT and Atg11-HA-SmBiT, or wild-type cells expressing Atg32 and Atg11 (negative control, N.C.), were grown in glycerol medium (Gly), collected at the OD600 = 1.4 point, incubated with substrates, and subjected to the NanoBiT-based bioluminescence assay. Data represent the averages of all experiments, with bars indicating standard deviations (*n* = 4 independent cultures). **(D)** Wild-type, *get1*Δ, *get2*Δ and *atg32*Δ cells expressing mito-DHFR-mCherry and wild-type *FAR9* or *FAR9-TM^ER^* were grown in glycerol medium (Gly), collected at the indicated time points, and subjected to western blotting. **(E)** The amounts of free mCherry in cells analyzed in **(D)** were quantified in three experiments. The signal intensity value of free mCherry in wild-type cells at the 72 h time point was set to 100%. Data represent the averages of all experiments, with bars indicating standard deviations (*n* = 3 independent cultures). Data were analyzed by two-tailed Student’s *t* test **(B, C, E)**.

### Artificial mitochondrial anchoring of the Far complex partially reduces mitophagy

As excess accumulation of the Far complex on the mitochondrial surface by loss of Get1/2 seems to perturb mitophagy, we sought to test if artificial targeting of the Far complex to mitochondria may suppress mitophagy without disrupting Get1/2 functions. The TA domains of Far9 and Far10 were replaced with those derived from Gem1 (Frederick et al., 2004), an OMM protein (Far9/Far10-TA^MITO^). We confirmed that Far8-3×GFP almost exclusively localizes to mitochondria in cells expressing Far9/Far10-TA^MITO^ (Fig. 6, A and B). In these cells, mitophagy under respiratory conditions was partially reduced (70% compared to wild-type cells) (Fig. 6, C and D). Importantly, this reduction was mostly abrogated in cells expressing Ppg1^H111N^, a catalytically inactive mutant (Fig. S5, A and B), suggesting that mitophagy suppression by the OMM-anchored Far complex requires Ppg1 phosphatase activity.

**Figure 6.**
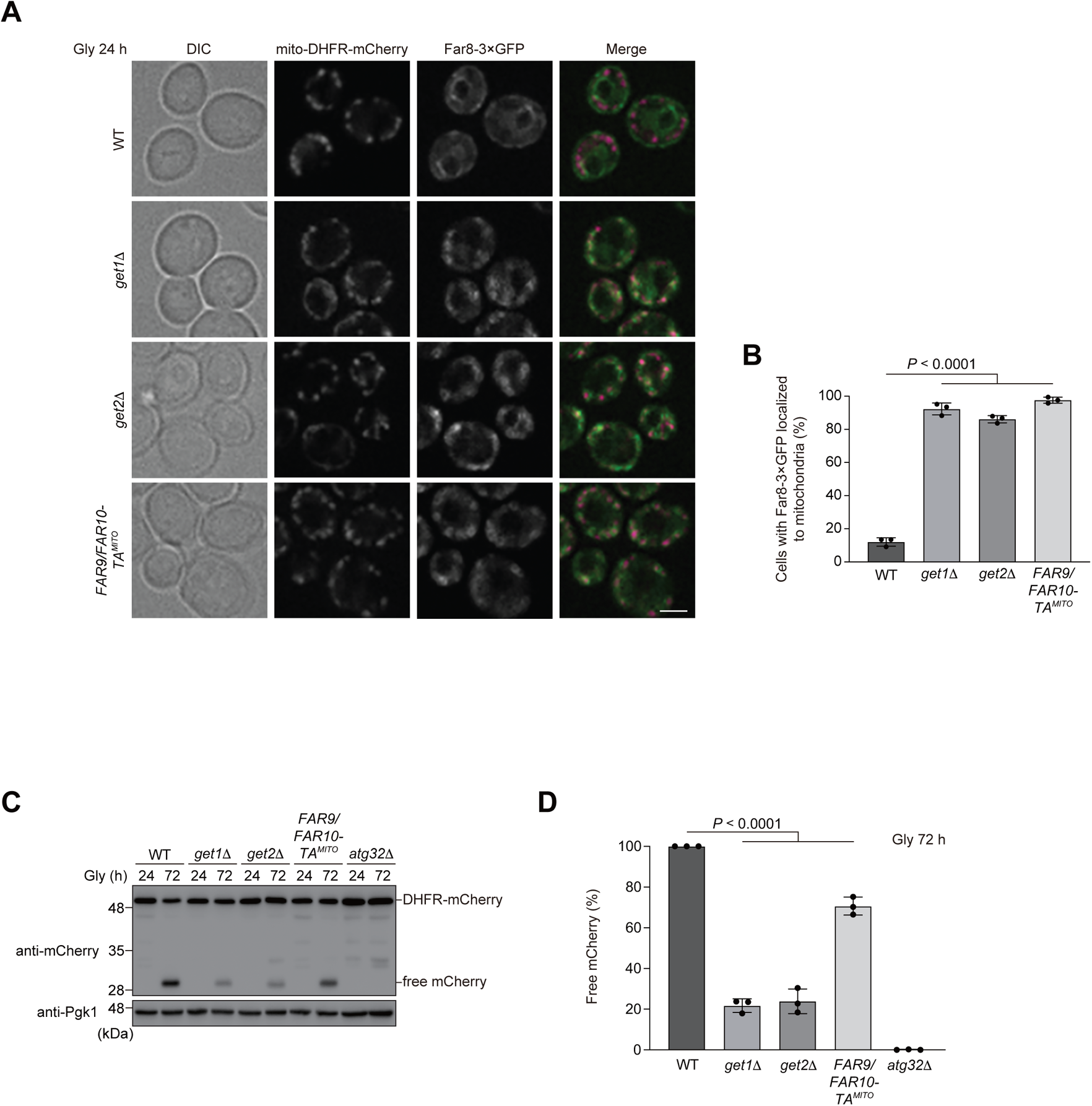
Artificial mitochondrial anchoring of the Far complex partially reduces mitophagy. **(A)** Representative images of wild-type, *get1*Δ, and *get2*Δ cells expressing the endogenous Far9/10 or a variant whose TA domains were replaced with the Gem1 TM domain (*FAR9/FAR10-TM^MITO^*) grown for 24 h in glycerol medium (Gly) and observed by structured illumination microscopy. All strains were derivatives expressing mito-DHFR-mCherry and Far8-3×GFP. Scale bar, 2 μm. **(B)** Cells analyzed in **(A)** were quantified in three experiments. Data represent the averages of all experiments, with bars indicating standard deviations (*n* = 3 independent cultures). **(C)** Wild-type, *get1*Δ, *get2*Δ, and *atg32*Δ cells expressing mito-DHFR-mCherry and the endogenous Far9/10 or Far9/Far10-TA^MITO^ were grown in glycerol medium (Gly), collected at the indicated time points, and subjected to western blotting. **(D)** The amounts of free mCherry in cells analyzed in **(C)** were quantified in three experiments. The signal intensity value of free mCherry in wild-type cells at the 72 h time point was set to 100%. Data represent the averages of all experiments, with bars indicating standard deviations (*n* = 3 independent cultures). Data were analyzed by one-way ANOVA with Dunnett’s multiple comparison test **(B, D)**.

### Msp1 is required for efficient mitophagy in cells lacking Get3

Previous studies demonstrate that Msp1, an OMM-anchored AAA-ATPase acting as an extractase, is important to remove non-mitochondrial TA proteins from the surface of mitochondria in the absence of Get components (Chen et al., 2014; Okreglak and Walter, 2014; Wang et al., 2020). Accordingly, we asked whether loss of Msp1 exacerbates mitophagy deficiencies in cells lacking the GET pathway. As double knockout of Msp1 and Get1/2 elicited extremely severe growth defects under respiratory conditions, we performed fluorescence microscopy and mitophagy assays for *msp1/get3*-double-null cells that could grow slowly with relatively mild phenotypes in liquid non-fermentable medium. Single knockout of Msp1 and Get3 slightly affected mitophagy (85% and 93%, respectively, compared to wild-type cells), whereas loss of these two proteins significantly compromised mitophagy (48% compared to wild-type cells) (Fig. 7, A and B). In addition, loss of Get3 in cells expressing Msp1^E193Q^ (Msp1EQ, an ATPase-inactive mutant) also synergistically disturbed degradation of mitochondria (51% compared to wild-type cells) (Fig. 7, A and B). These results suggest that Msp1 ATPase activity is critical to prevent mitophagy suppression in GET-deficient cells.

**Figure 7.**
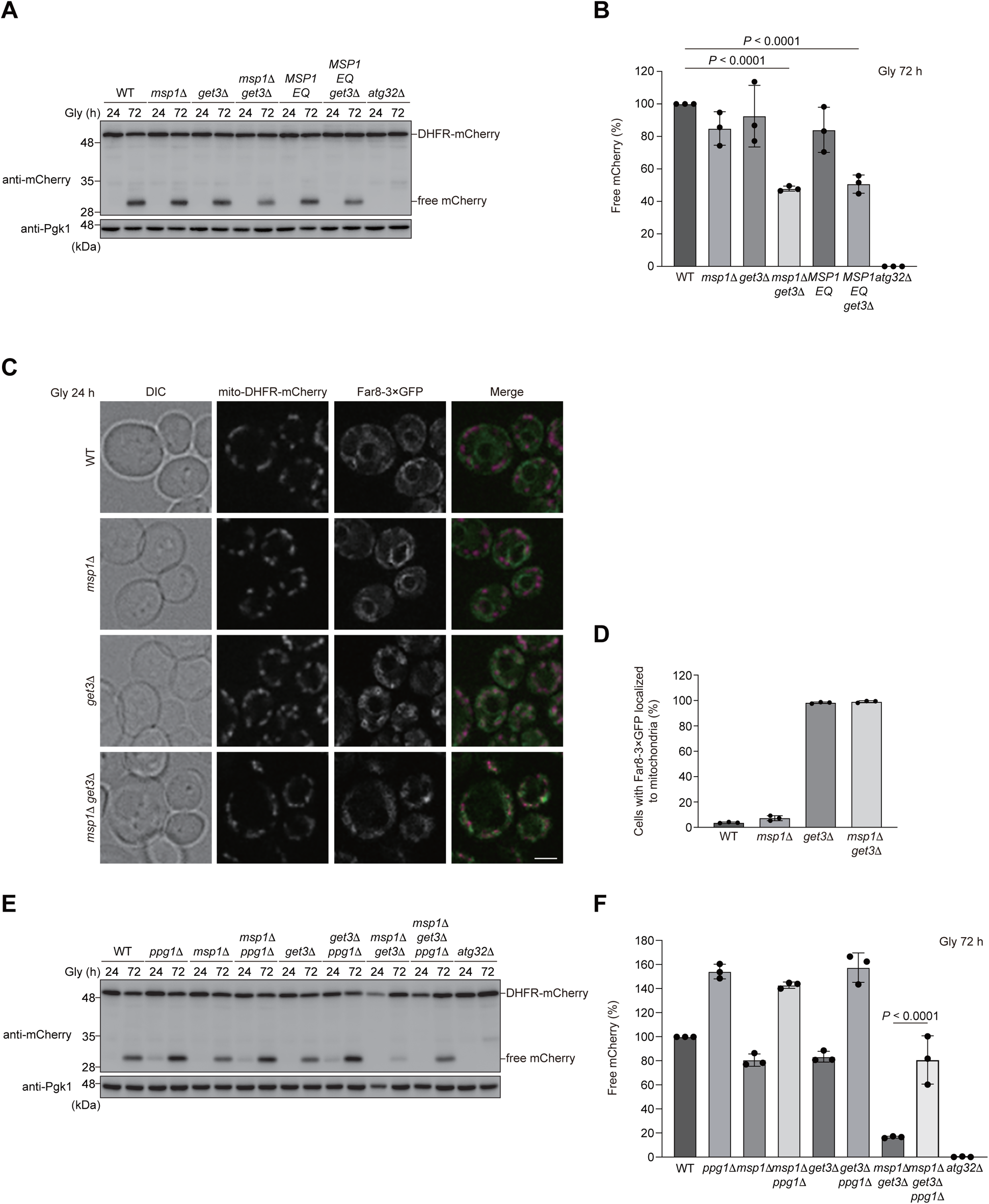
Msp1 is required for efficient mitophagy in cells lacking Get3. **(A)** Wild-type, *msp1*Δ, *get3*Δ, *msp1*Δ *get3*Δ, the endogenous *MSP1*-expressing or *MSP1 E193Q* (*MSP1EQ*)-expressing *get3*Δ, and *atg32*Δ cells were grown in glycerol medium (Gly), collected at the indicated time points, and subjected to western blotting. All strains were derivatives expressing mito-DHFR-mCherry. **(B)** The amount of free mCherry in cells analyzed in **(A)** was quantified in three experiments. The signal intensity value of free mCherry in wild-type cells at the 72 h time point was set to 100%. Data represent the averages of all experiments, with bars indicating standard deviations (*n* = 3 independent cultures). **(C)** Representative images of wild-type, *msp1*Δ, *get3*Δ, and *msp1*Δ *get3*Δ cells expressing mito-DHFR-mCherry and Far8-3×GFP grown for 24 h in glycerol medium (Gly) and observed by structured illumination microscopy. Scale bar, 2 μm. **(D)** Cells analyzed in **(C)** were quantified in three experiments. Data represent the averages of all experiments, with bars indicating standard deviations (*n* = 3 independent cultures). **(E)** Wild-type, *ppg1*Δ, *msp1*Δ, *get3*Δ, *msp1*Δ *ppg1*Δ, *get3*Δ *ppg1*Δ, *msp1*Δ *get3*Δ, *msp1*Δ *get3*Δ *ppg1*Δ, and *atg32*Δ cells expressing mito-DHFR-mCherry were grown in glycerol medium (Gly), collected at the indicated time points, and subjected to western blotting. **(F)** The amount of free mCherry in cells analyzed in **(E)** was quantified in three experiments. The signal intensity value of free mCherry in wild-type cells at the 72 h time point was set to 100%. Data represent the averages of all experiments, with bars indicating standard deviations (*n* = 3 independent cultures). Data were analyzed by one-way ANOVA with Dunnett’s multiple comparison test **(B)**, or two-tailed Student’s *t* test **(F)**.

Next, we performed fluorescence microscopy and found that loss of Msp1 did not significantly affect ER localization of Far8-3×GFP (Fig. 7, C and D). By contrast, Far8-3×GFP localized to mitochondria in *get3*-null and *msp1/get3*-double-null cells (Fig. 7, C and D). Based on these observations, we investigated if loss of Ppg1 affects mitophagy in *msp1*/*get3*-double-null cells, and found that *msp1/get3/ppg1*-triple-null cells significantly restored mitophagy (81% compared to wild-type cells) (Fig. 7, E and F). Similarly, expression of Atg32(Δ151-200), a deletion mutant lacking a domain required for Ppg1-mediated dephosphorylation, also increased mitophagy in cells lacking Get3 and Msp1 (60% compared to wild-type cells) (Fig. S5, C and D). Collectively, these results support the idea that the GET pathway and Msp1 cooperatively act to prevent suppression of mitophagy by the Ppg1-Far complex.

## Discussion

In the present study, we show that the GET pathway contributes to Atg32 phosphorylation by promoting localization of the Ppg1-Far phosphatase complex to the ER (Fig. 8). Loss of Get1/2 (ER membrane-anchored insertase), or Get3 (cytosolic ATPase), partially reduces Atg32 phosphorylation, thereby abrogating Atg32-Atg11 interactions in the early phase of respiratory growth (Fig. 1, A and C). Consistent with this observation, mitophagy is severely compromised in *get1/2*-null cells under prolonged respiration (Onishi et al., 2018). However, cells lacking Get3 exhibit only minor defects in mitophagy (Onishi et al., 2018), raising the possibility that in the prolonged phase of respiratory growth, Get1/2 may have unappreciated additional function(s) to promote mitophagy independently of its insertase activity (Fig. S2, B-D), or that unknown protein(s) may exert a Get3-related compensatory role in promoting mitophagy.

**Figure 8.**
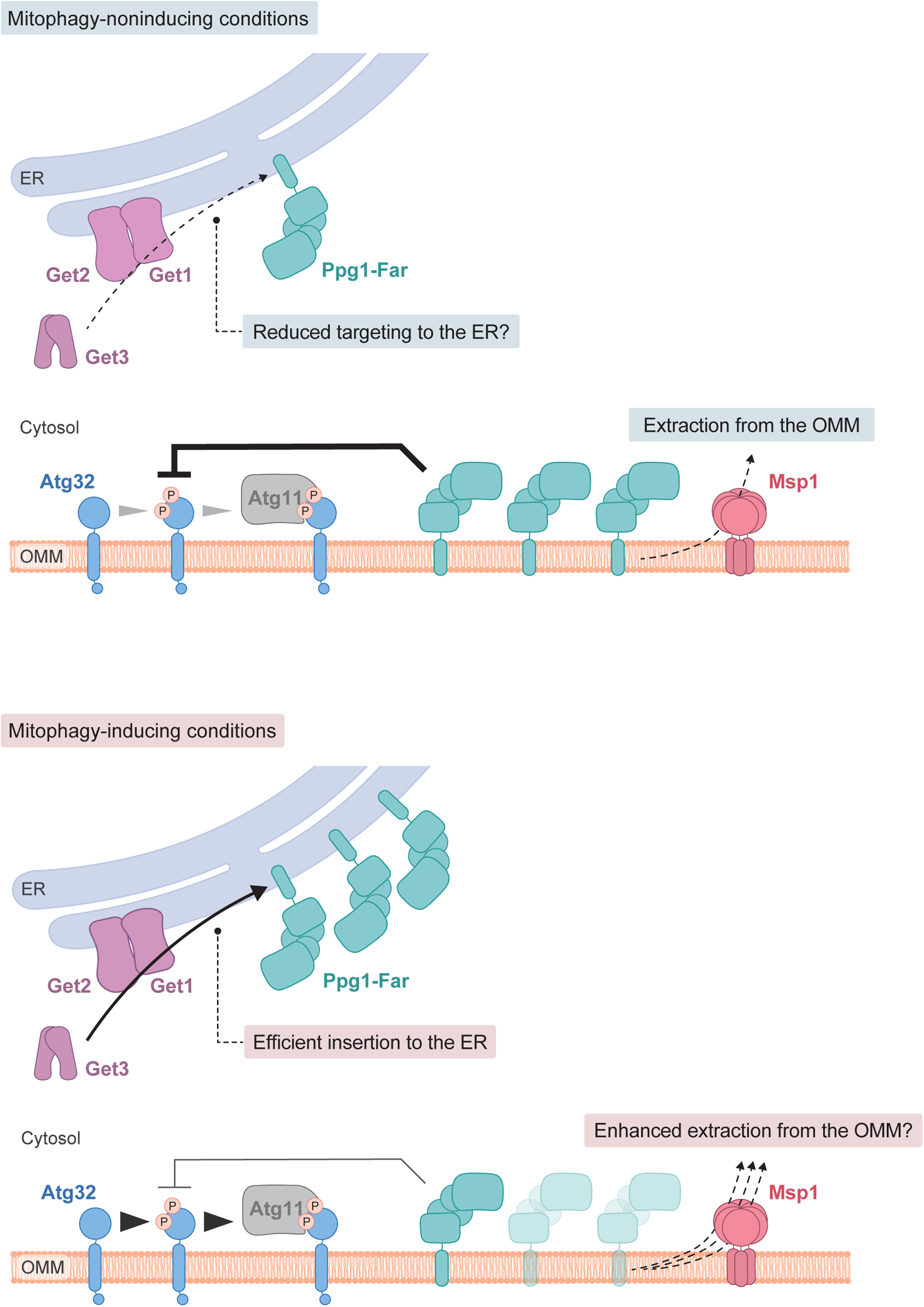
A hypothetical model for activation of Atg32-mediated mitophagy. **(Upper panel)** Under mitophagy-noninducing (fermentable) conditions, a substantial fraction of the Ppg1-Far complex escapes the GET pathway that may have reduced levels, activity, and/or affinity, localizes to mitochondria, and suppress Atg32 phosphorylation and Atg32-Atg11 interactions. Mitochondria-anchored Ppg1-Far can be extracted from the OMM via Msp1. **(Lower panel)** Under mitophagy-inducing (non-fermentable) conditions, the GET pathway efficiently mediates targeting of the Ppg1-Far complex to the ER, which in turn promotes Atg32 phosphorylation and Atg32-Atg11 interactions. Msp1-dependent extraction of mitochondria-anchored Ppg1-Far from the OMM may be enhanced by unknown mechanisms, contributing to activation of mitophagy.

Evidently, Atg32-Atg11 interactions and mitophagy in cells lacking Get1/2 can mostly be restored by additional loss of Ppg1, a phosphatase that dephosphorylates Atg32 (Fig. 2, A-F), suggesting that Ppg1 is likely to be the primary cause of reduced Atg32 phosphorylation in *get1/2*-null mutants. Consistent with these findings, loss of Far9, a component of the Far complex that binds to Ppg1 and acts in a cooperative manner to dephosphorylate Atg32, also increased Atg32-Atg11 interactions and mitophagy in Get1/2-deficient cells (Fig. 4, A-C). Far9 is an ER-resident TA protein of the Far complex (Pracheil and Liu, 2013), and loss of the Get1/2 insertase activity perturbs ER localization of the Far complex (Fig. 3, A-D; and Fig. S2, A and B), supporting the idea that the GET pathway promotes insertion of the Far TA proteins to the ER membrane.

Although disruption of the GET pathway leads to targeting of multiple ER-resident TA proteins to mitochondria (Jonikas et al., 2009; Schuldiner et al., 2008), how these ectopically targeted proteins impact events on the mitochondrial surface remains enigmatic. Upon loss of Get components, the Far complex predominantly targets to mitochondria (Fig. 3, A-D; and Fig. S2, A and B) in a manner dependent on the TA proteins Far9 and Far10 (Fig. 3, E and F). Anchoring to the mitochondrial surface seems to be critical for the Far complex to efficiently abrogate Atg32-Atg11 interactions and mitophagy, since cytosolic diffusion or GET-independent ER anchoring of the Far complex leads to restoration of those processes in *get1/2*-null cells (Fig. 4, A-C; and Fig. S3, A-C; and Fig. 5, C-E). Based on the observations from us (Fig. S2, E and F) and others (Innokentev et al., 2020) that a fraction of the Far complex localizes to mitochondria even in wild-type cells, we favor a hypothetical model that dynamic changes in the GET pathway (*e.g.* expression level, insertase activity, and substrate affinity) could affect the number of Ppg1-Far complex targeted to the ER or mitochondria, thereby serving as a regulatory process for mitophagy (Fig. 8). Further studies are needed to test this hypothesis.

During the course of this study, we noticed that our several results seem to be somewhat different from the recently reported data on localization and function of the Ppg1-Far complex (Innokentev et al., 2020). First, we demonstrate that the Far complex is mostly localized to the ER in cells during non-fermentable growth (Fig. 3 A-D), whereas it has been shown that the Far complex is distributed almost equally to both mitochondria and the ER in cells during fermentable growth (Innokentev et al., 2020). These distinct features might be due to different growth condition (mitophagy-inducing or -noninducing). Second, cells containing the GET-independently ER-localized Far complex exhibit mitophagy at near wild-type levels under prolonged respiration (Fig. 5, D and E), whereas cells containing the Far9-Cyb5^TA^-dependently ER-localized Far complex have been shown to accelerate mitophagy at the early stationary phase (Innokentev et al., 2020). These differences might result from TM segments (one derived from the non-TA protein Sec12 or TA protein Cyb5) and/or mitophagy assay time points (72 h or 40 h in non-fermentable medium). Third, we show that Gem1 TA-dependent artificial targeting of the Far complex to mitochondria causes a partial defect in mitophagy under prolonged respiration (Fig. 6, C and D), whereas it has been demonstrated that mitochondria-targeted Far complex by Tom5 TA strongly diminishes mitophagy at the early stationary phase (Innokentev et al., 2020). This phenotypic difference might be attributed to TA domains used for mitochondrial anchoring and/or mitophagy assay time points (72 h or 40 h in non-fermentable medium). Nevertheless, it seems possible that the mitochondria-anchored Ppg1-Far complex could suppress stationary-phase mitophagy more effectively at the early phase than the late phase.

Expression of the Get1/2 insertase-inactive mutants leads to extensive accumulation of the Ppg1-Far complex on the mitochondrial surface, while mitophagy is only partially decreased in these mutant cells (70% compared to wild-type cells) (Fig. S2, B-D). Notably, these phenotypes are similar to those in *get3*-null cells (Fig. 7, A-D) (Onishi et al., 2018), which is in agreement with the previous finding that the Get1/2 insertase-inactive mutants cannot recruit Get3 to the ER (Wang et al., 2011). In addition, artificial targeting of the Ppg1-Far complex to mitochondria only partially reduces mitophagy under prolonged respiration (70% compared to wild-type cells) (Fig. 6, C and D). Together, these findings raise the possibility that Get1/2 may be a bifunctional complex acting as a general insertase for ER-resident TA proteins, and serving as a pro-mitophagic factor independently of its insertase activity.

Finally, our data reveal a potential role of the OMM-anchored AAA-ATPase Msp1 in mitophagy. Consistent with the previous reports that Msp1 extracts non-mitochondrial TA proteins from the mitochondrial surface upon loss of Get components (Chen et al., 2014; Okreglak and Walter, 2014; Wang et al., 2020; Wohlever et al., 2017), cells lacking both Get3 and Msp1 display synthetic defects in mitophagy that can be rescued by loss of Ppg1 or expression of an Atg32 variant defective for interaction with the Ppg1-Far complex (Fig. 7, A and B, E and F; and Fig. S5, C and D). Thus, although ER localization of the Far complex seems to be hardly altered in cells lacking Msp1 (Fig. 7, C and D), it remains possible that this OMM-anchored extractase may act in removal of non-mitochondrial TA proteins, such as Far9 and Far10, thereby contributing to Atg32 phosphorylation, Atg32-Atg11 interactions, and mitophagy (Fig. 8). How the GET pathway and Msp1 coordinately act in activation of Atg32-mediated mitophagy awaits further investigations.

## Materials and methods

### Yeast strains and plasmids used in this study

Yeast strains and plasmids used in this thesis are listed in Table S1 and S2. Standard genetic and molecular biology methods were performed for generating yeast strains.

### Growth conditions of yeast

Yeast cells were incubated in YPD medium (1% yeast extract, 2% peptone and 2% dextrose), synthetic medium (0.17% yeast nitrogen base without amino acids and ammonium sulfate, 0.5% ammonium sulfate) with 0.5% casamino acids and either 2% dextrose (SDCA), or 0.1% dextrose plus 3% glycerol (SDGCA), supplemented with the necessary amino acids. For mitophagy assay under respiratory conditions, cells grown to mid-log phase in SDCA were transferred to SDGCA and incubated at 30°C.

### Protein phosphatase treatment assays

For protein phosphatase assays, cells were pre-grown in SDCA, and transferred to SDGCA. 2.0 OD600 units of cells were collected and subjected to alkaline lysis and TCA (Trichloroacetic acid) precipitation. The pellet was resuspended in a reaction buffer (50 mM Tris-HCl pH 7.5, 100 mM NaCl, 2 mM DTT, 0.5 mM EDTA, 0.01% Brij-35, 2 mM MgCl2), treated with or without lambda protein phosphatase (λ-PPase) in the presence or absence of PPase inhibitor at 30 °C for 1 h. Samples corresponding to 0.2 OD600 units of cells were loaded per lane

### Structured illumination microscopy

Live yeast cells expressing Far8-3×GFP were observed using a structured illumination microscopy (Stefer et al.). Differential interference contrast (DIC) and fluorescence images were obtained under a KEYENCE BZ-X810 system equipped with a 100× objective lens (CFI Apochromat TIRF 100XC Oil, Plan-APO TIRF 100, NA: 1.49; Nikon), filter sets for GFP and mCherry (BZ-X filter GFP and BZ-X filter TRITC, respectively; KEYENCE). Cell images were captured using acquisition and analysis software (BZ-X800 Analyzer; KEYENCE).

### Western blotting

Samples corresponding to 0.1-0.4 OD600 units of cells were separated by SDS-PAGE followed by western blotting and immunodecoration with primary antibodies raised against mCherry (1:2,000, Abcam ab125096), Pgk1 (1:10,000, Abcam, ab113687), GFP (1:1000, Roche, 13921700), HA (1:5,000, Sigma, A2095). After treatment with the secondary antibodies, horseradish peroxidase (HRP)-conjugated rabbit anti-mouse IgG (H + L) for mCherry, GFP, HA, Pgk1, followed by the enhanced chemiluminescence reagent Western Lightning Plus-ECL (PerkinElmer, 203-19151) or ImmunoStar LD (Wako, PTJ2005), proteins were detected using a luminescent image analyzer (FUSION Solo S; VILBER). Quantification of the signals was performed using FUSION Solo S (VILBER).

### Bioluminescence assay for protein-protein interactions

For quantitative analysis of Atg32-Atg11 interactions using NanoLuc Binary Technology (NanoBiT, Promega), Atg32 fused to 3 copies of GFP and Large BiT (LgBiT; 17.6 kDa), and Atg11 fused to Small BiT (SmBiT; 11 amino acids) were expressed endogenously (constructed by Yang Liu, Osaka University, Japan). Upon interaction of Atg32 and Atg11 with each other, SmBiT and LgBiT are brought into close proximity, leading to structural complementation and generation of a luminescent signal. For the assay, cells were grown in glycerol media (SDGCA). 1.0 OD600 units of cells were collected in the early phase of respiration (OD600: 1.4~1.6) and washed with 400 µl PBS. After washing, cells were dissolved in 40 µl PBS and applied to a 96 well plate. The detection reagent was prepared by diluting the Nano-Glo Live Cell Substrate (Promega, 0000360026) with the Nano-Glo LCS Dilution Buffer (Promega, 0000333050) to make the Nano-Glo Live Cell Reagent. 10 µl diluted detection reagent was added onto the 96 well plate and mixed with the cells. Then, cells were incubated at 37°C for 1 hour. After incubation, the luminescent signal was detected by the microplate reader (Fluoroskan Ascent FL; Thermo Fisher Scientific) (exposure time: 1,000 ms). For the detection of the GFP fluorescent signal derived from Atg32, 1.0 OD600 units of cells were collected at the same time point, and dissolved in 100 µl SDGCA media, applied to the 96 well plate. GFP signal was measured by microplate reader (Fluoroskan Ascent FL; Thermo Fisher Scientific) (excitation: 485 nm, emission: 538 nm, exposure time: 1,000 ms), and used to normalize the luminescence intensity.

### Statistical analysis

Results are presented as means including ± standard deviation. Statistical analyses were performed with Excel for Mac (Microsoft) and GraphPad Prism 9 (GraphPad Software), using two-tailed Student’s *t*-test and one-way ANOVA followed by Tukey’s or Dunnett’s multiple comparison test. All the statistical tests performed are indicated in the figure legends.

### Online supplemental material

Fig. S1 shows a schematic description on the NanoBiT assays for Atg32-Atg11 interactions, and the results on Atg32-Atg11 interactions and mitophagy in *get1/2*-null cells expressing a catalytically inactive Ppg1 mutant. Fig. S2 contains the data from microscopic imaging, mitophagy assay, and processing assay for cells expressing Far8-3×GFP and mito-DHFR-mCherry. Fig. S3 shows the results on Atg32-Atg11 interactions and mitophagy in *get1/2*-null cells lacking Far10. Fig. S4 contains the data from microscopic imaging for *get1/2*-null cells expressing an artificially ER-anchored Far9. Fig. S5 shows the results on mitophagy in cells expressing a catalytically inactive Ppg1 mutant and an ectopically mitochondria-targeted Far9/10. Table S1 contains a list of yeast strains used in this study. Table S2 shows a list of plasmid used in this study.

## Acknowledgements

We thank Miyuki Sato (Gunma University, Japan) for valuable suggestions on artificial ER anchoring, Elmar Schiebel (Heidelberg University, Germany) for kindly providing us with the plasmid pFA6a-3myeGFP-kanMX6, and Yang Liu (Osaka University, Japan) for providing us with the NanoBiT assay strains. This work was supported in part by JSPS KAKENHI Grants JP19J10384, JP21K15041 (to M.O.), JP16H04784, JP19H03222, and JP20H05324 (to KO), and the Osaka University International Joint Research Promotion Programs (Type A+ and Type A-GKP) (to KO). The authors declare no competing financial interests.

## Author contributions

M. Onishi and K. Okamoto obtained funding and conceptualized the study. M. Onishi and K. Okamoto designed experiments. M. Onishi performed experiments. M. Onishi and K. Okamoto wrote the manuscript.

## Figure legends

**Figure S1.**
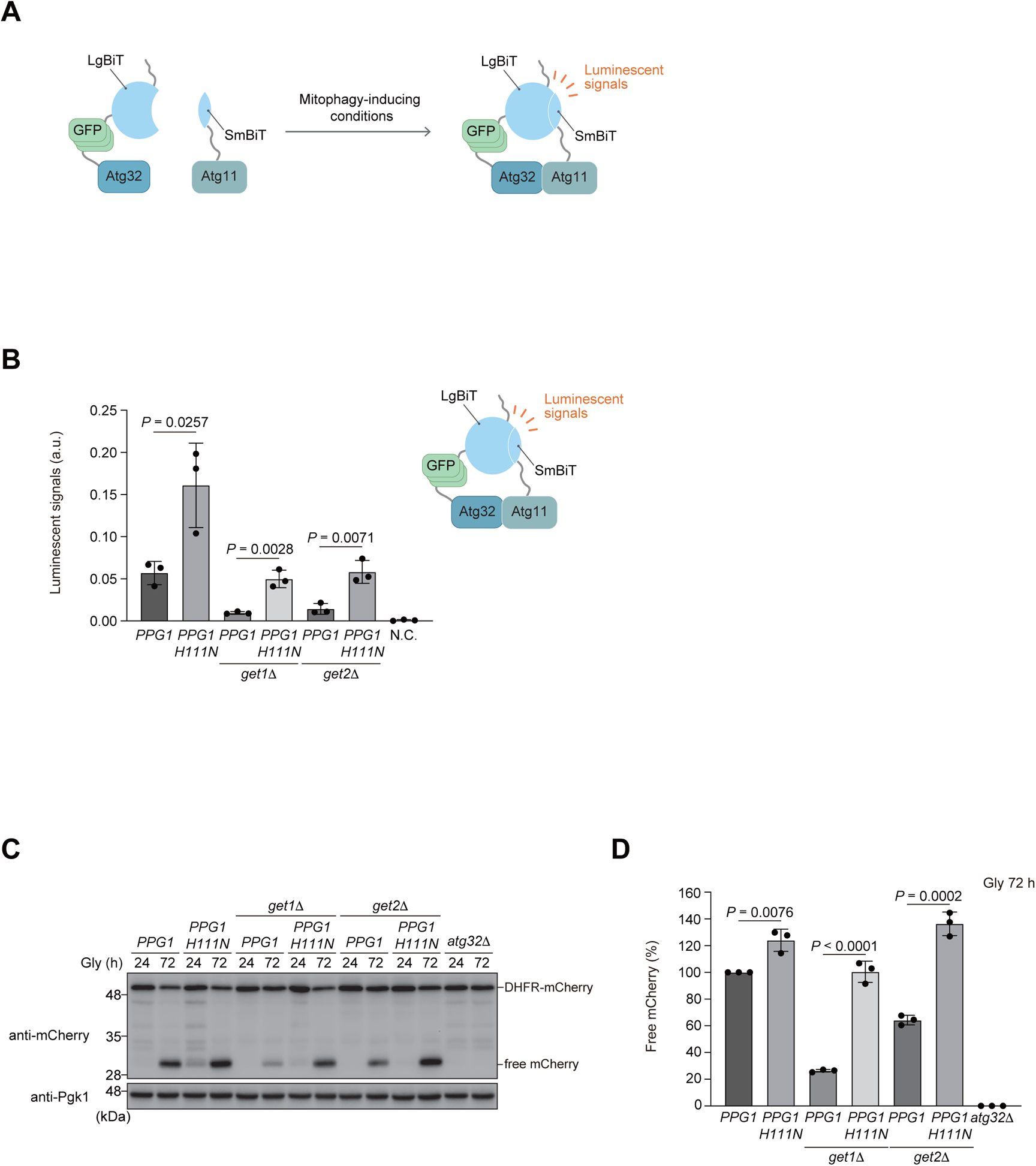
Expression of a catalytically inactive Ppg1 mutant restores Atg32-Atg11 interactions and mitophagy in *get1/2*-null cells. **(A)** A schematic illustration of the NanoBiT system using cells expressing Atg32-3HA-3×GFP-3FLAG-LgBiT and Atg11-HA-SmBiT. Upon mitophagy induction, Atg32 interacts with Atg11, bringing each luminescent subunit into close proximity to form a functional unit that releases a luminescent signal, which is detected by a microplate reader. **(B)** Derivatives of cells analyzed in **(C)** expressing Atg32-3HA-3×GFP-3FLAG-LgBiT and Atg11-HA-SmBiT, or wild-type cells expressing Atg32 and Atg11 (negative control, N.C.), were grown in glycerol medium (Gly), collected at the OD600 = 1.4 point, incubated with substrates, and subjected to the NanoBiT-based bioluminescence assay. Data represent the averages of all experiments, with bars indicating standard deviations (*n* = 3 independent cultures). **(C)** Wild-type, *get1*Δ, and *get2*Δ cells containing chromosomally integrated wild-type *PPG1* or *PPG1 H111N* were grown in glycerol medium (Gly), collected at the indicated time points, and subjected to western blotting. **(D)** The amounts of free mCherry in cells analyzed in **(C)** was quantified in three experiments. The signal intensity value of free mCherry in wild-type cells at the 72 h time point was set to 100%. Data represent the averages of all experiments, with bars indicating standard deviations (*n* = 3 independent cultures). Data were analyzed by two-tailed Student’s *t* test **(B, D)**.

**Figure S2.**
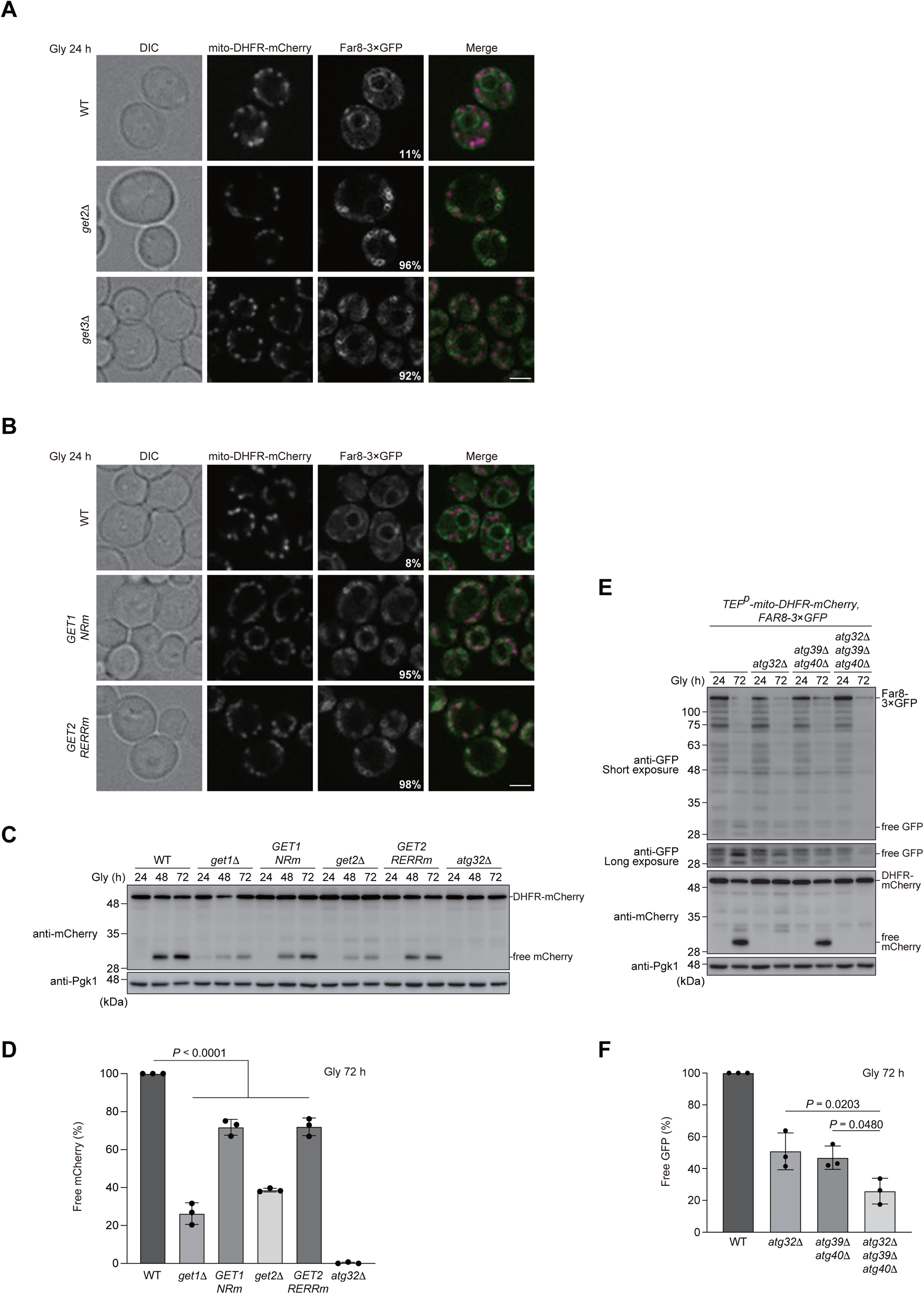
ER localization of the Far complex is perturbed in GET-deficient cells. **(A)** Representative images of wild-type, *get2*Δ, and *get3*Δ cells expressing mito-DHFR-mCherry and Far8-3×GFP grown for 24 h in glycerol medium (Gly) and observed by structured illumination microscopy. Scale bar, 2 μm. The percentage of cells exhibiting mitochondria-localized Far8-3×GFP is depicted. **(B)** Representative images of wild-type cells expressing the endogenous *GET1/2*, chromosomally integrated *GET1 NRm (N72A, R73A*), or a *GET2 RERRm (R14E, E15R, R16E, R17E*) grown for 24 h in glycerol medium (Gly) and observed by structured illumination microscopy. All strains were derivatives expressing mito-DHFR-mCherry and Far8-3×GFP. Scale bar, 2 µm. The percentage of cells exhibiting mitochondria-localized Far8-3×GFP is depicted. **(C)** Wild-type, *get1*Δ, *get2*Δ, or cells expressing chromosomally integrated *GET1 NRm* or *GET2 RERRm,* and *atg32*Δ cells were grown in glycerol medium (Gly), collected at the indicated time points, and subjected to western blotting. **(D)** The amounts of free mCherry in cells analyzed in **(C)** was quantified in three experiments. The signal intensity value of free mCherry in wild-type cells at the 72 h time point was set to 100%. Data represent the averages of all experiments, with bars indicating standard deviations (*n* = 3 independent cultures). **(E)** Wild-type, *atg32*Δ, *atg39*Δ *atg40*Δ, and *atg32*Δ *atg39*Δ *atg40*Δ cells expressing mito-DHFR-mCherry and Far8-3×GFP were grown in glycerol medium (Gly), collected at the indicated time points, and subjected to western blotting. Generation of free GFP indicates transport of Far8 to the vacuole. **(F)** The amount of free GFP in cells analyzed in **(E)** was quantified in three experiments. The signal intensity value of free GFP in wild-type cells at the 72 h time point was set to 100%. Data represent the averages of all experiments, with bars indicating standard deviations (*n* = 3 independent cultures). Data were analyzed by one-way ANOVA with Dunnett’s multiple comparison test **(D)** or Tukey’s multiple comparison test **(F)**.

**Figure S3.**
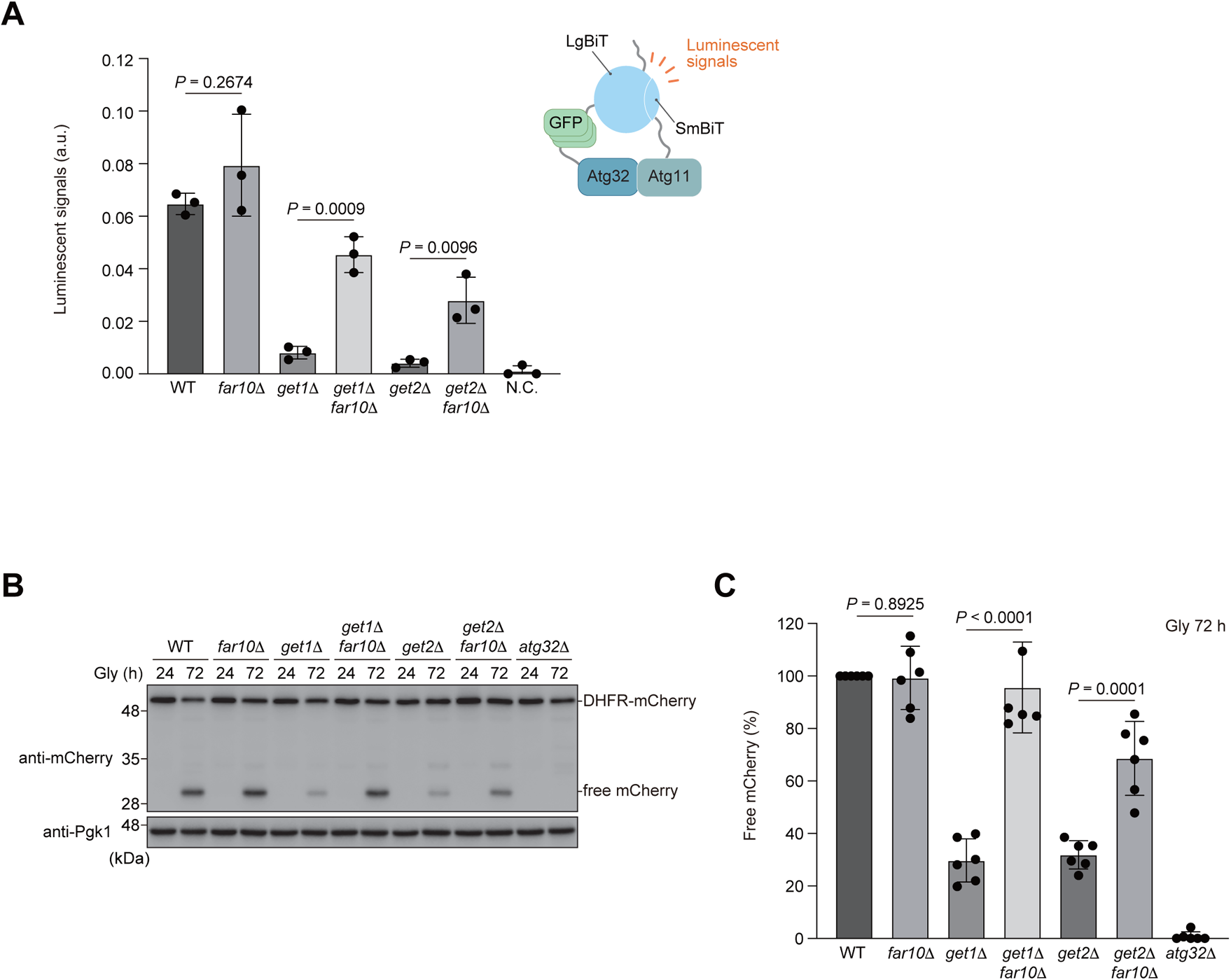
Loss of Far10 significantly ameliorates mitophagic deficiencies in the absence of Get1/2. **(A)** Wild-type, *far10*Δ, *get1*Δ, *get2*Δ, *get1*Δ *far10*Δ, and *get2*Δ *far10*Δ cells expressing Atg32-3HA-3×GFP-3FLAG-LgBiT and Atg11-HA-SmBiT, or wild-type cells expressing Atg32 and Atg11 (negative control, N.C.) were grown in glycerol medium (Gly), collected at the OD600 = 1.4 point, incubated with substrates, and subjected to the NanoBiT-based bioluminescence assay. Data represent the averages of all experiments, with bars indicating standard deviations (*n* = 3 independent cultures). **(B)** Wild-type, *far10*Δ, *get1*Δ, *get2*Δ, *get1*Δ *far10*Δ, *get2*Δ *far10*Δ, and *atg32*Δ cells expressing mito-DHFR-mCherry were grown in glycerol medium (Gly), collected at the indicated time points, and subjected to western blotting. **(C)** The amount of free mCherry in cells analyzed in **(B)** was quantified in six experiments. The signal intensity value of free mCherry in wild-type cells at the 72 h time point was set to 100%. Data represent the averages of all experiments, with bars indicating standard deviations (*n* = 6 independent cultures). Data were analyzed by two-tailed Student’s *t* test **(A, C)**.

**Figure S4.**
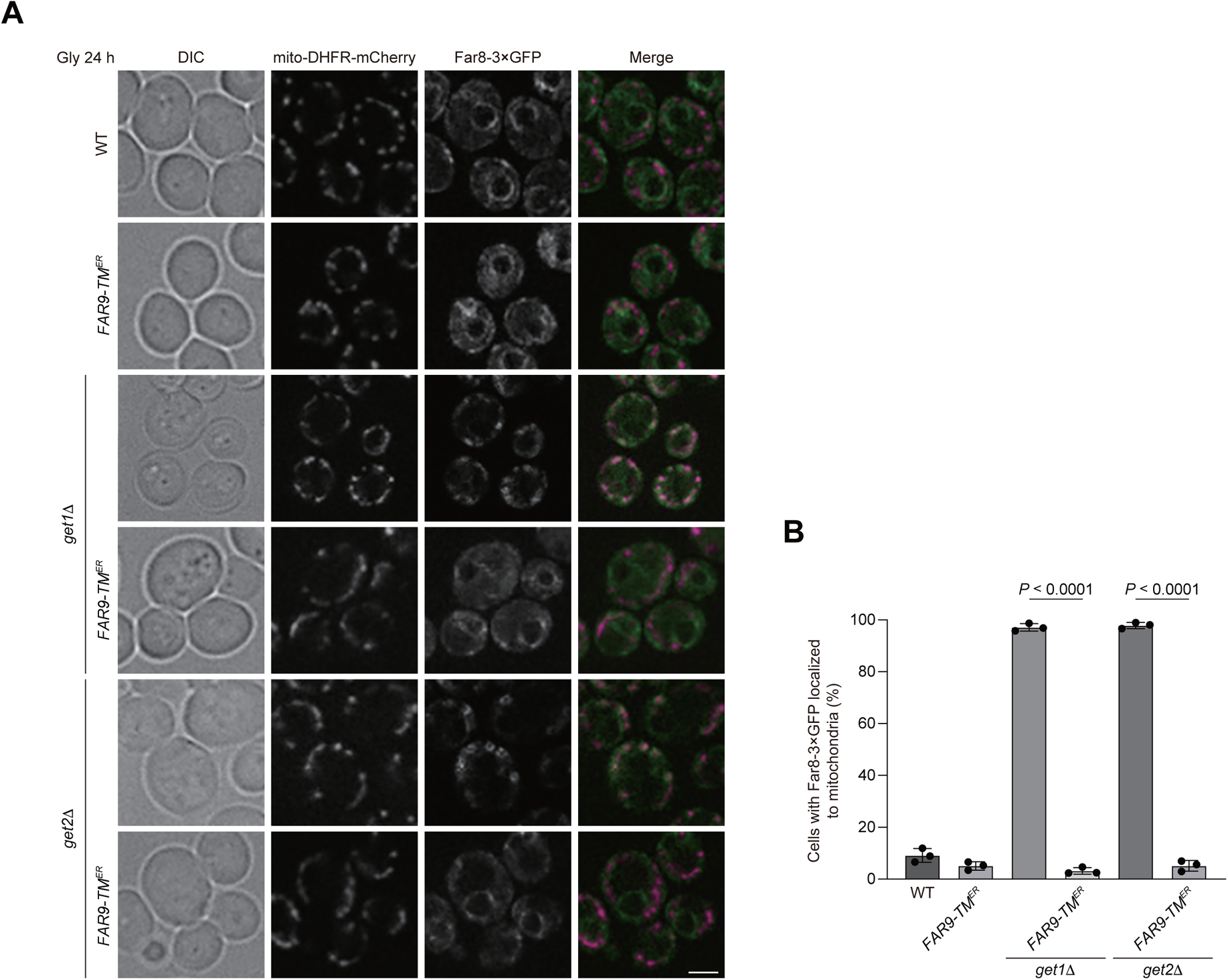
Artificial ER anchoring of the Far complex in *get1/2*-null cells. **(A)** Representative images of wild-type, *get1*Δ, and *get2*Δ cells expressing the endogenous Far9 or a variant whose TA domain was replaced with the Sec12 TM domain (*FAR9-TM^ER^*) grown for 24 h in glycerol medium (Gly) and observed by structured illumination microscopy. All strains were derivatives expressing mito-DHFR-mCherry and Far8-3×GFP. Scale bar, 2 μm. **(B)** Cells analyzed in **(A)** were quantified in three experiments. Data represent the averages of all experiments, with bars indicating standard deviations. Data were analyzed by two-tailed Student’s *t* test **(B)**.

**Figure S5.**
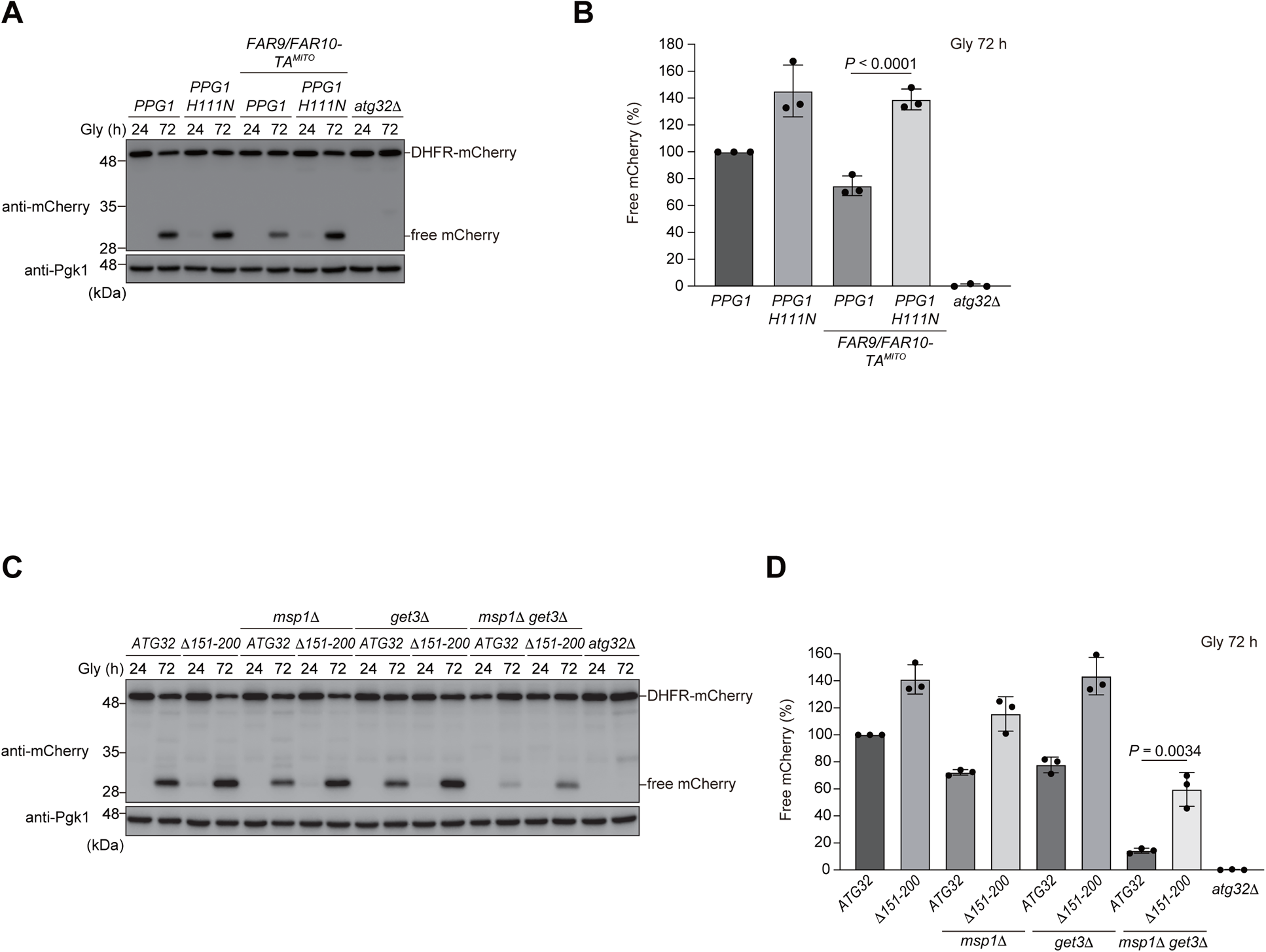
Suppression of mitophagy by artificially mitochondria-anchored Ppg1-Far and restoration of mitophagy in *get3/msp1*-double-null cells expressing a Atg32 variant defective in Ppg1-Far interaction. **(A)** Wild-type cells expressing the endogenous *FAR9* and *PPG1* or chromosomally integrated *FAR9/FAR10-TA^MITO^* and *PPG1 H111N* were grown in glycerol medium (Gly), collected at the indicated time points, and subjected to western blotting. **(B)** The amounts of free mCherry in cells analyzed in **(A)** were quantified in three experiments. The signal intensity value of free mCherry in wild-type cells at the 72 h time point was set to 100%. Data represent the averages of all experiments, with bars indicating standard deviations (*n* = 3 independent cultures). Data were analyzed by two-tailed Student’s *t* test **(B)**. **(C)** Wild-type, *msp1*Δ, *get3*Δ, and *msp1*Δ *get3*Δ cells expressing the endogenous *ATG32* or chromosomally integrated *ATG32(Δ151-200)* were grown in glycerol medium (Gly), collected at the indicated time points, and subjected to western blotting. **(D)** The amounts of free mCherry in cells analyzed in **(C)** were quantified in three experiments. The signal intensity value of free mCherry in wild-type cells at the 72 h time point was set to 100%. Data represent the averages of all experiments, with bars indicating standard deviations (*n* = 3 independent cultures). Data were analyzed by two-tailed Student’s *t* test **(B and D)**.

**Table S1.**
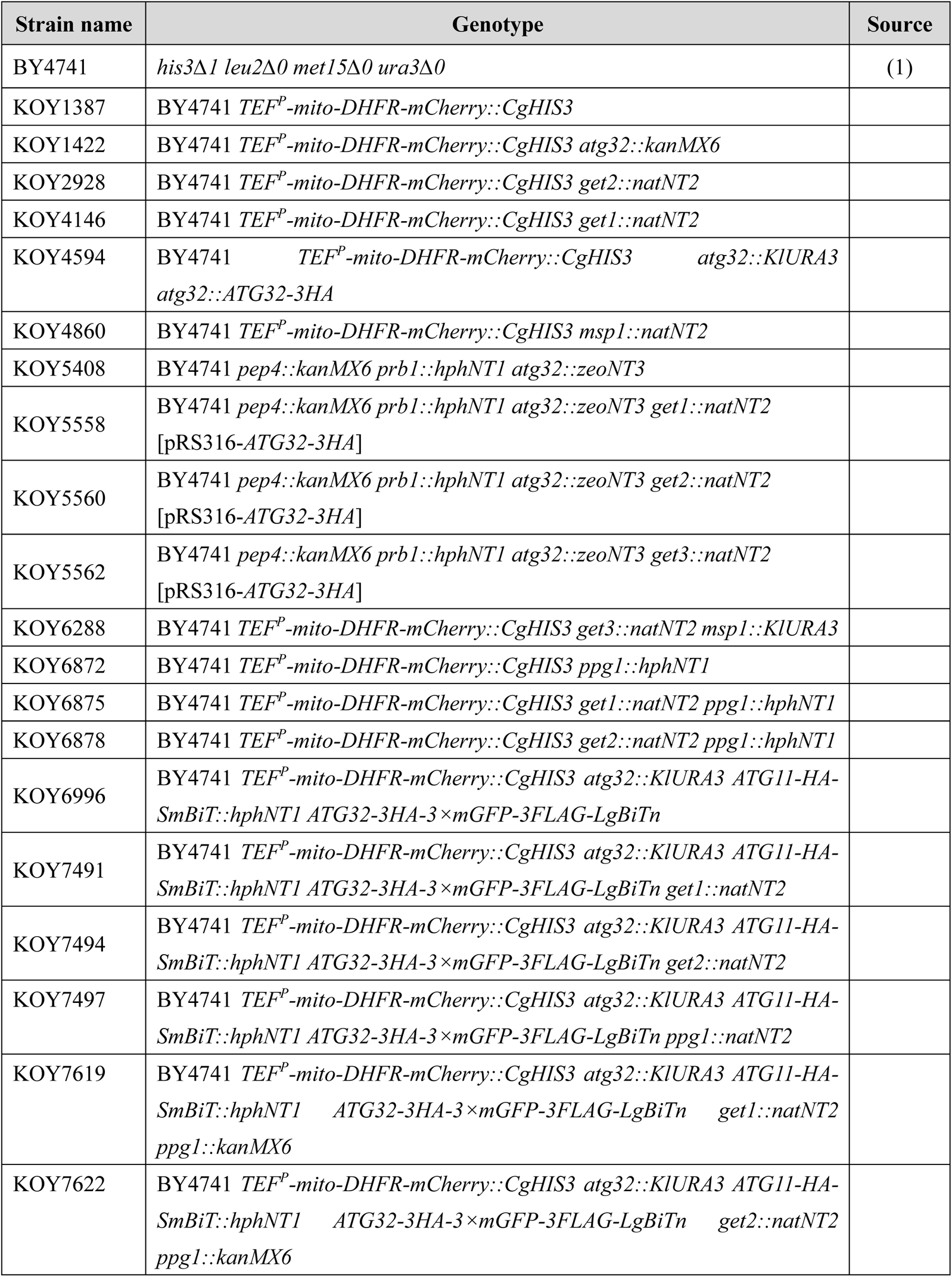
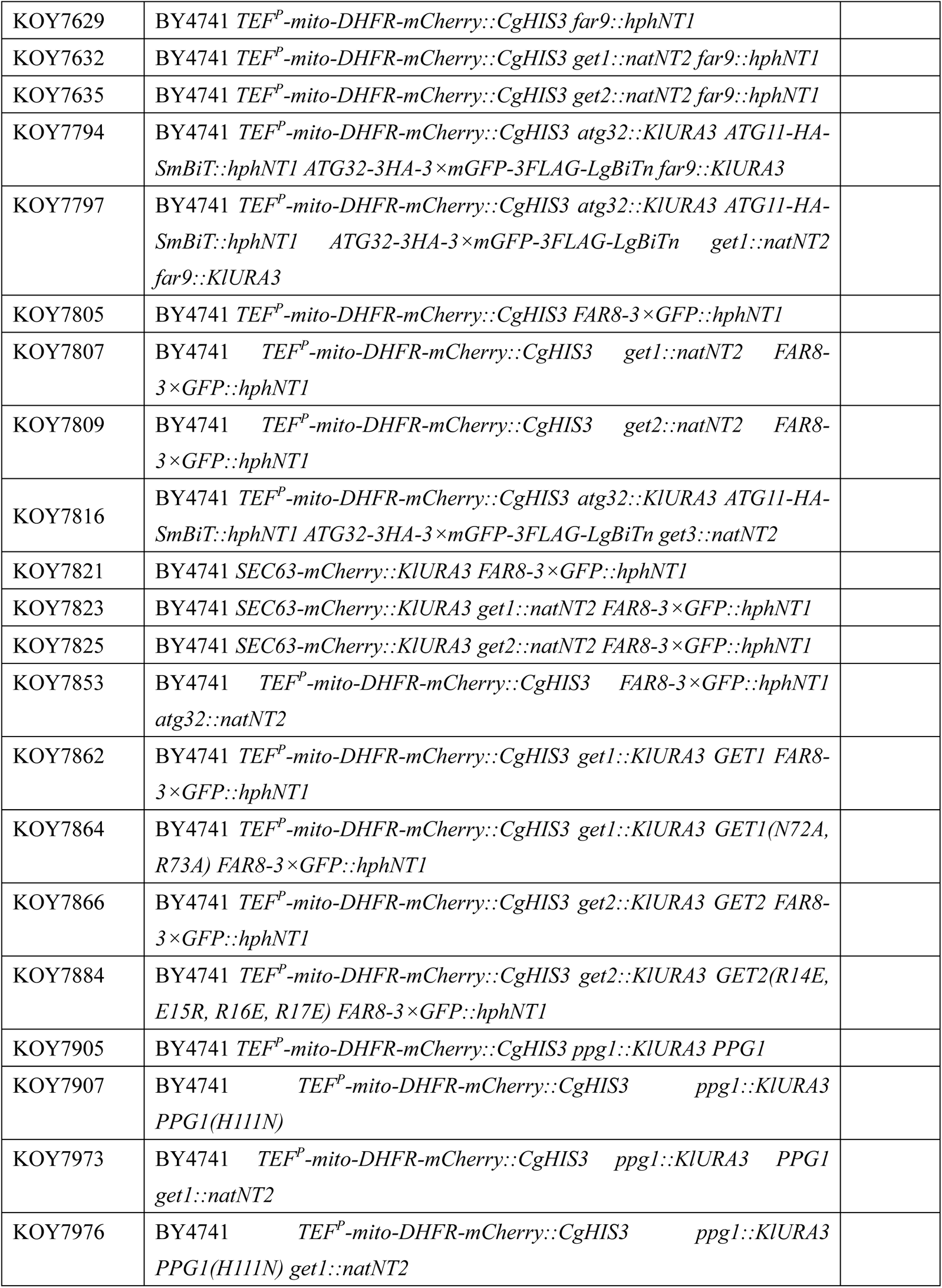
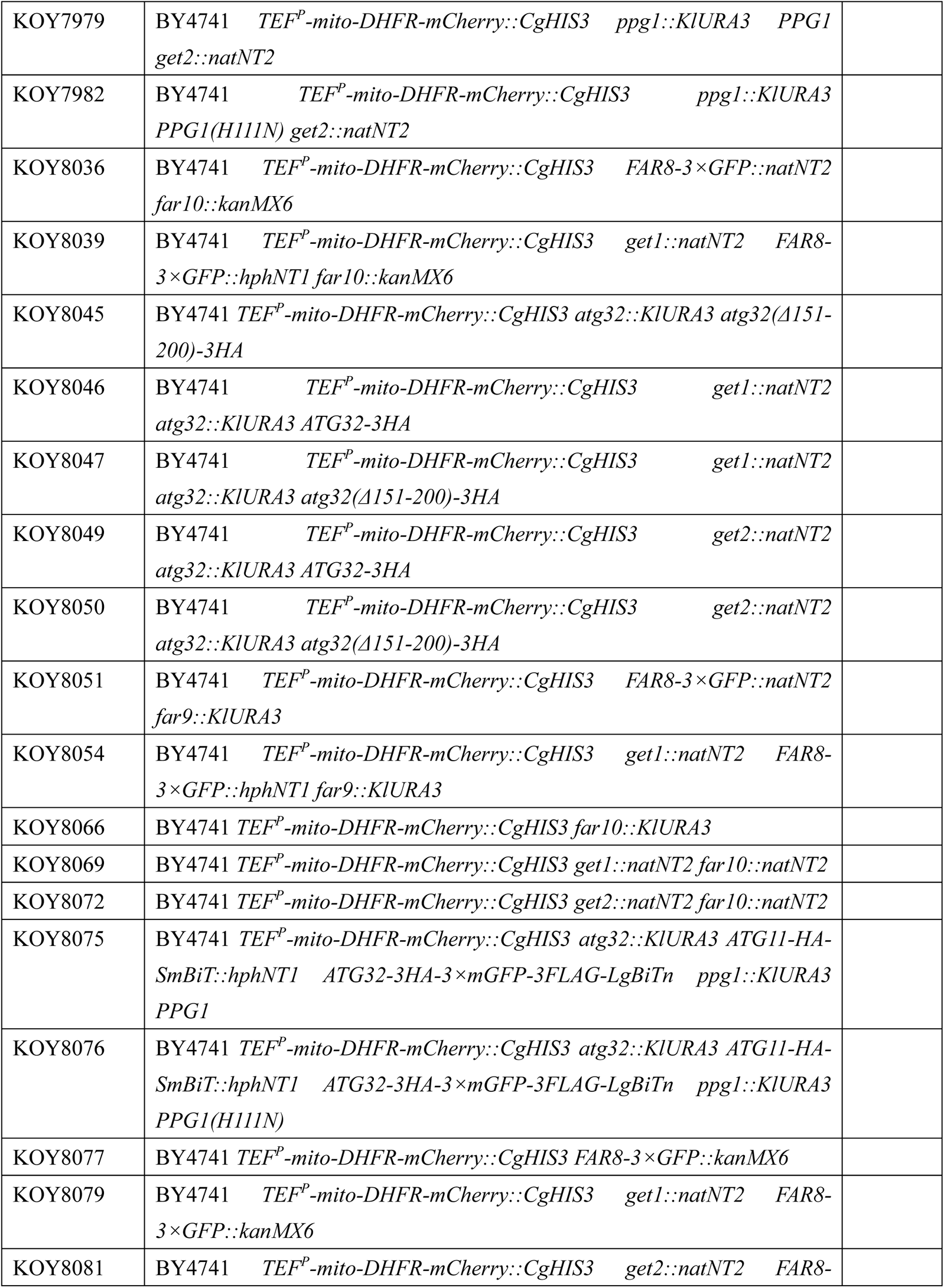
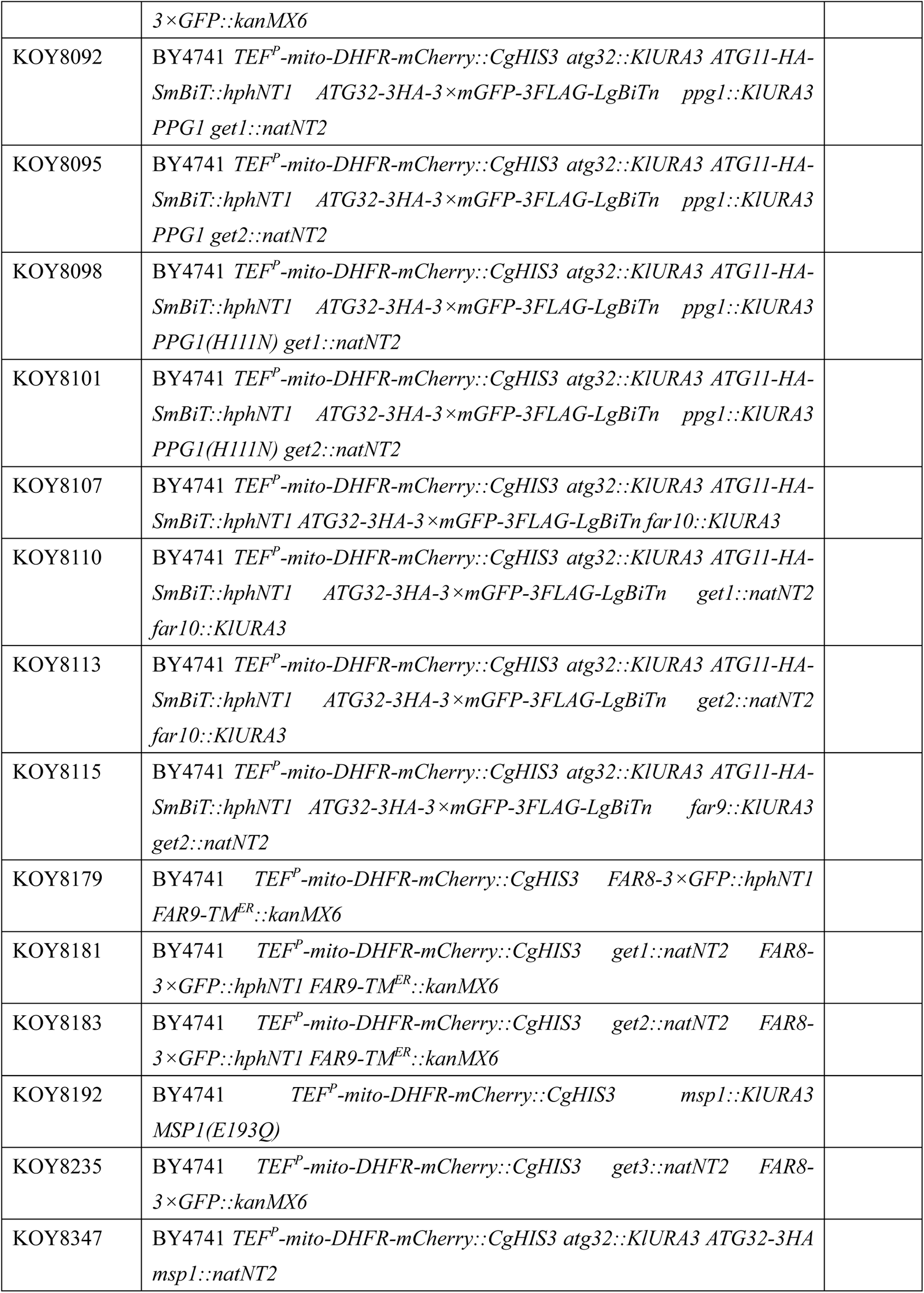
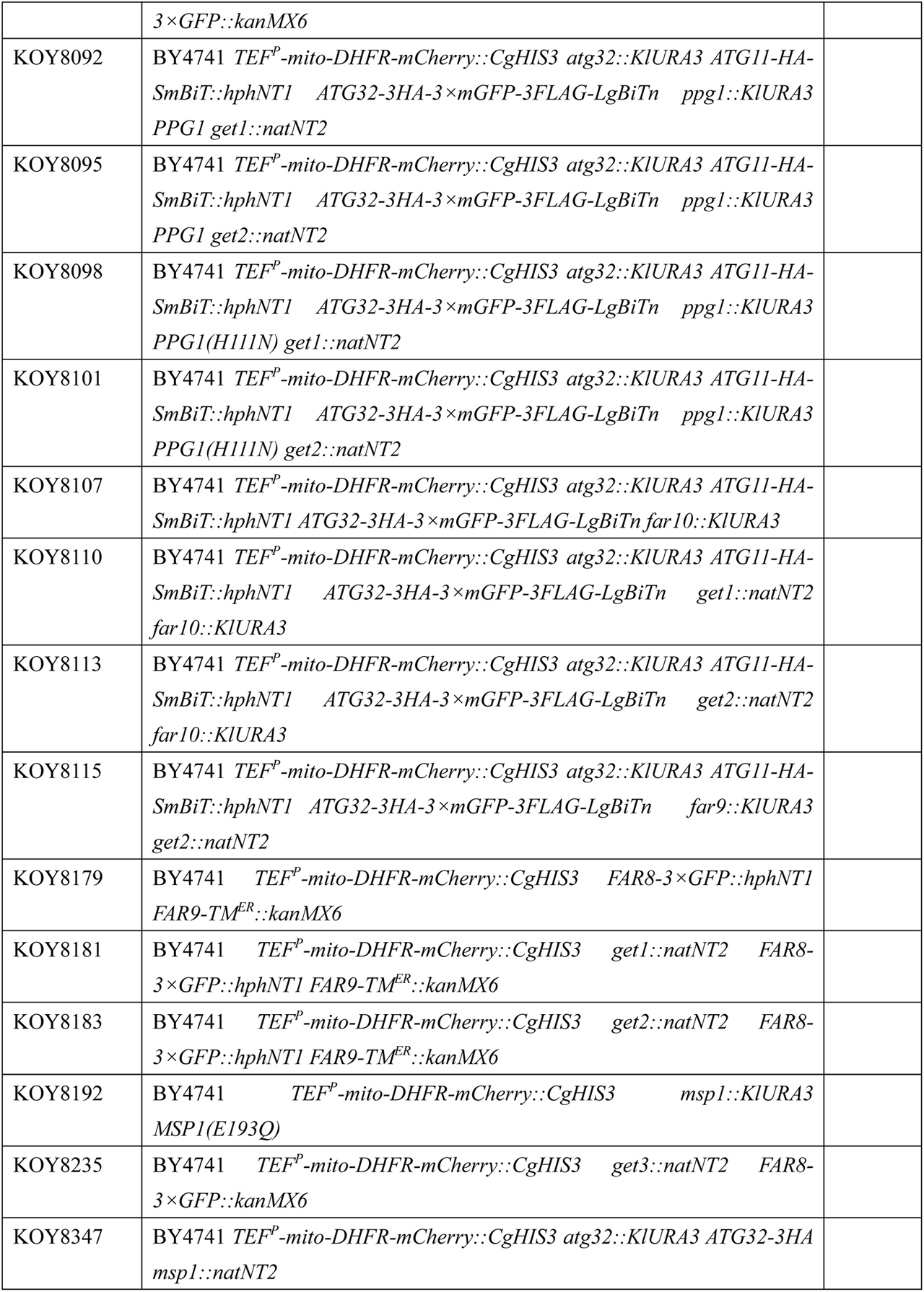
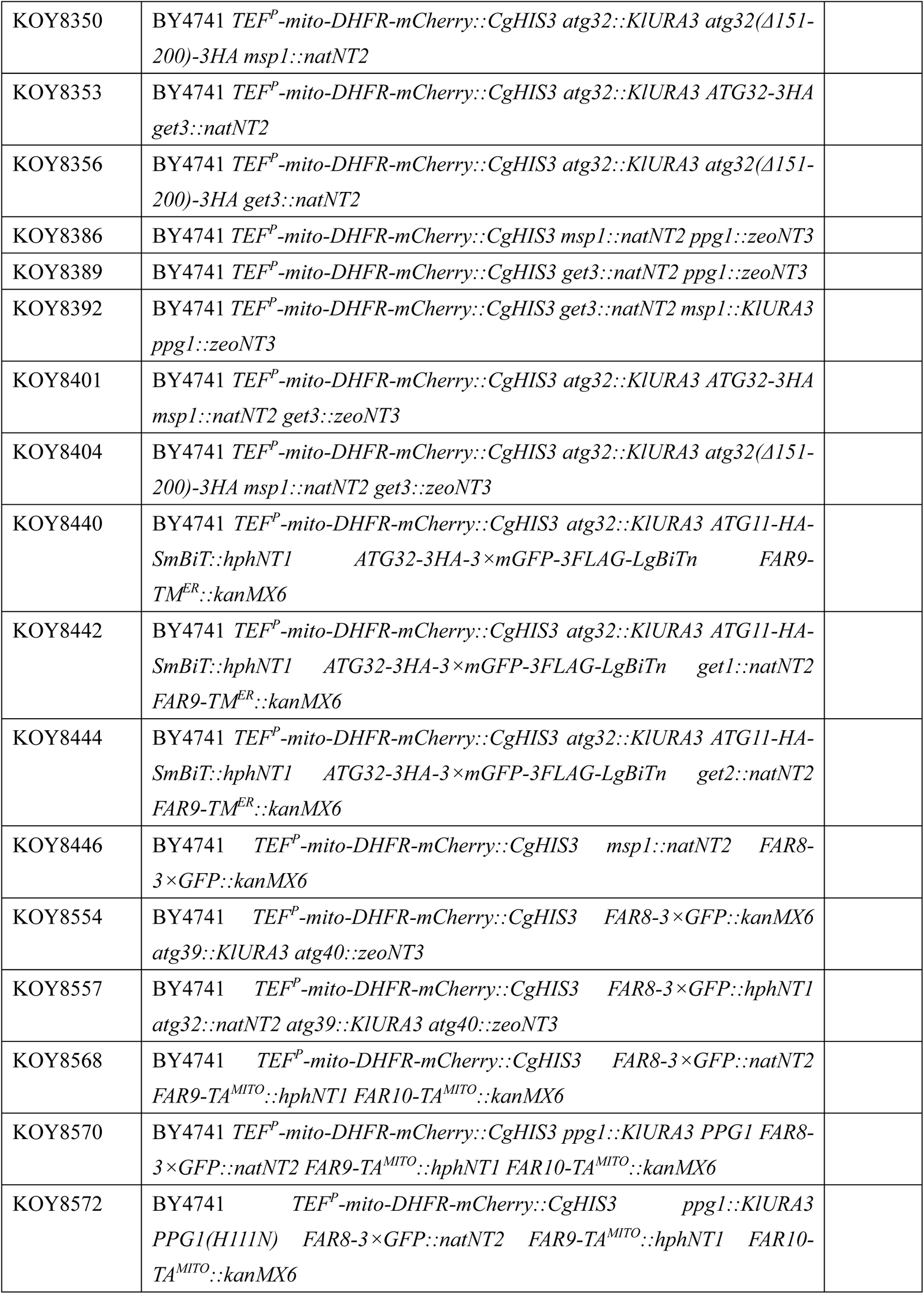
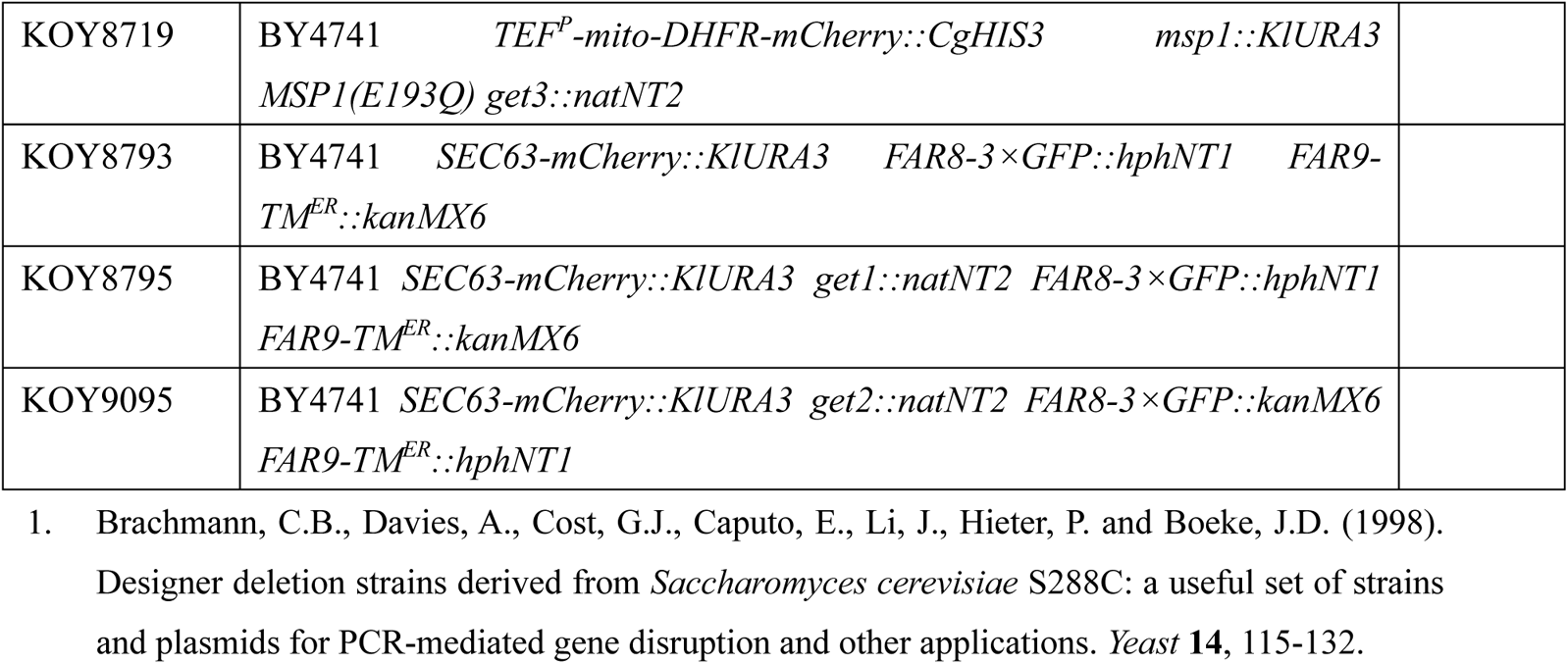
Yeast strains used in this study.

| Strain name | Genotype | Source |
| --- | --- | --- |
| BY4741 | *his3Δ1 leu2Δ0 met15Δ0 ura3Δ0* | (1) |
| KOY1387 | BY4741 *TEF<sup>P</sup>-mito-DHFR-mCherry::CgHIS3* |  |
| KOY1422 | BY4741 *TEF<sup>P</sup>-mito-DHFR-mCherry::CgHIS3 atg32::kanMX6* |  |
| KOY2928 | BY4741 *TEF<sup>P</sup>-mito-DHFR-mCherry::CgHIS3 get2::natNT2* |  |
| KOY4146 | BY4741 *TEF<sup>P</sup>-mito-DHFR-mCherry::CgHIS3 get1::natNT2* |  |
| KOY4594 | BY4741 *TEF<sup>P</sup>-mito-DHFR-mCherry::CgHIS3 atg32::KIURA3 atg32::ATG32-3HA* |  |
| KOY4860 | BY4741 *TEF<sup>P</sup>-mito-DHFR-mCherry::CgHIS3 msp1::natNT2* |  |
| KOY5408 | BY4741 *pep4::kanMX6 prb1::hphNT1 atg32::zeoNT3* |  |
| KOY5558 | BY4741 *pep4::kanMX6 prb1::hphNT1 atg32::zeoNT3 get1::natNT2 [pRS316-ATG32-3HA]* |  |
| KOY5560 | BY4741 *pep4::kanMX6 prb1::hphNT1 atg32::zeoNT3 get2::natNT2 [pRS316-ATG32-3HA]* |  |
| KOY5562 | BY4741 *pep4::kanMX6 prb1::hphNT1 atg32::zeoNT3 get3::natNT2 [pRS316-ATG32-3HA]* |  |
| KOY6288 | BY4741 *TEF<sup>P</sup>-mito-DHFR-mCherry::CgHIS3 get3::natNT2 msp1::KIURA3* |  |
| KOY6872 | BY4741 *TEF<sup>P</sup>-mito-DHFR-mCherry::CgHIS3 ppg1::hphNT1* |  |
| KOY6875 | BY4741 *TEF<sup>P</sup>-mito-DHFR-mCherry::CgHIS3 get1::natNT2 ppg1::hphNT1* |  |
| KOY6878 | BY4741 *TEF<sup>P</sup>-mito-DHFR-mCherry::CgHIS3 get2::natNT2 ppg1::hphNT1* |  |
| KOY6996 | BY4741 *TEF<sup>P</sup>-mito-DHFR-mCherry::CgHIS3 atg32::KIURA3 ATG11-HA-SmBiT::hphNT1 ATG32-3HA-3×mGFP-3FLAG-LgBiTn* |  |
| KOY7491 | BY4741 *TEF<sup>P</sup>-mito-DHFR-mCherry::CgHIS3 atg32::KIURA3 ATG11-HA-SmBiT::hphNT1 ATG32-3HA-3×mGFP-3FLAG-LgBiTn get1::natNT2* |  |
| KOY7494 | BY4741 *TEF<sup>P</sup>-mito-DHFR-mCherry::CgHIS3 atg32::KIURA3 ATG11-HA-SmBiT::hphNT1 ATG32-3HA-3×mGFP-3FLAG-LgBiTn get2::natNT2* |  |
| KOY7497 | BY4741 *TEF<sup>P</sup>-mito-DHFR-mCherry::CgHIS3 atg32::KIURA3 ATG11-HA-SmBiT::hphNT1 ATG32-3HA-3×mGFP-3FLAG-LgBiTn ppg1::natNT2* |  |
| KOY7619 | BY4741 *TEF<sup>P</sup>-mito-DHFR-mCherry::CgHIS3 atg32::KIURA3 ATG11-HA-SmBiT::hphNT1 ATG32-3HA-3×mGFP-3FLAG-LgBiTn get1::natNT2 ppg1::kanMX6* |  |
| KOY7622 | BY4741 *TEF<sup>P</sup>-mito-DHFR-mCherry::CgHIS3 atg32::KIURA3 ATG11-HA-SmBiT::hphNT1 ATG32-3HA-3×mGFP-3FLAG-LgBiTn get2::natNT2 ppg1::kanMX6* |  |
| KOY7629 | BY4741 *TEF<sup>P</sup>-mito-DHFR-mCherry::CgHIS3 far9::hphNT1* |  |
| KOY7632 | BY4741 *TEF<sup>P</sup>-mito-DHFR-mCherry::CgHIS3 get1::natNT2 far9::hphNT1* |  |
| KOY7635 | BY4741 *TEF<sup>P</sup>-mito-DHFR-mCherry::CgHIS3 get2::natNT2 far9::hphNT1* |  |
| KOY7794 | BY4741 *TEF<sup>P</sup>-mito-DHFR-mCherry::CgHIS3 atg32::KIURA3 ATG11-HA-SmBiT::hphNT1 ATG32-3HA-3×mGFP-3FLAG-LgBiTn far9::KIURA3* |  |
| KOY7797 | BY4741 *TEF<sup>P</sup>-mito-DHFR-mCherry::CgHIS3 atg32::KIURA3 ATG11-HA-SmBiT::hphNT1 ATG32-3HA-3×mGFP-3FLAG-LgBiTn get1::natNT2 far9::KIURA3* |  |
| KOY7805 | BY4741 *TEF<sup>P</sup>-mito-DHFR-mCherry::CgHIS3 FAR8-3×GFP::hphNT1* |  |
| KOY7807 | BY4741 *TEF<sup>P</sup>-mito-DHFR-mCherry::CgHIS3 get1::natNT2 FAR8-3×GFP::hphNT1* |  |
| KOY7809 | BY4741 *TEF<sup>P</sup>-mito-DHFR-mCherry::CgHIS3 get2::natNT2 FAR8-3×GFP::hphNT1* |  |
| KOY7816 | BY4741 *TEF<sup>P</sup>-mito-DHFR-mCherry::CgHIS3 atg32::KIURA3 ATG11-HA-SmBiT::hphNT1 ATG32-3HA-3×mGFP-3FLAG-LgBiTn get3::natNT2* |  |
| KOY7821 | BY4741 *SEC63-mCherry::KIURA3 FAR8-3×GFP::hphNT1* |  |
| KOY7823 | BY4741 *SEC63-mCherry::KIURA3 get1::natNT2 FAR8-3×GFP::hphNT1* |  |
| KOY7825 | BY4741 *SEC63-mCherry::KIURA3 get2::natNT2 FAR8-3×GFP::hphNT1* |  |
| KOY7853 | BY4741 *TEF<sup>P</sup>-mito-DHFR-mCherry::CgHIS3 FAR8-3×GFP::hphNT1 atg32::natNT2* |  |
| KOY7862 | BY4741 *TEF<sup>P</sup>-mito-DHFR-mCherry::CgHIS3 get1::KIURA3 GET1 FAR8-3×GFP::hphNT1* |  |
| KOY7864 | BY4741 *TEF<sup>P</sup>-mito-DHFR-mCherry::CgHIS3 get1::KIURA3 GET1(N72A, R73A) FAR8-3×GFP::hphNT1* |  |
| KOY7866 | BY4741 *TEF<sup>P</sup>-mito-DHFR-mCherry::CgHIS3 get2::KIURA3 GET2 FAR8-3×GFP::hphNT1* |  |
| KOY7884 | BY4741 *TEF<sup>P</sup>-mito-DHFR-mCherry::CgHIS3 get2::KIURA3 GET2(R14E, E15R, R16E, R17E) FAR8-3×GFP::hphNT1* |  |
| KOY7905 | BY4741 *TEF<sup>P</sup>-mito-DHFR-mCherry::CgHIS3 ppg1::KIURA3 PPG1* |  |
| KOY7907 | BY4741 *TEF<sup>P</sup>-mito-DHFR-mCherry::CgHIS3 ppg1::KIURA3 PPG1(H111N)* |  |
| KOY7973 | BY4741 *TEF<sup>P</sup>-mito-DHFR-mCherry::CgHIS3 ppg1::KIURA3 PPG1 get1::natNT2* |  |
| KOY7976 | BY4741 *TEF<sup>P</sup>-mito-DHFR-mCherry::CgHIS3 ppg1::KIURA3 PPG1(H111N) get1::natNT2* |  |
| KOY7979 | BY4741 *TEF<sup>P</sup>-mito-DHFR-mCherry::CgHIS3* *ppg1::KIURA3* *PPG1*<br>*get2::natNT2* |  |
| KOY7982 | BY4741 *TEF<sup>P</sup>-mito-DHFR-mCherry::CgHIS3* *ppg1::KIURA3*<br>*PPG1(H111N)* *get2::natNT2* |  |
| KOY8036 | BY4741 *TEF<sup>P</sup>-mito-DHFR-mCherry::CgHIS3* *FAR8-3×GFP::natNT2*<br>*far10::kanMX6* |  |
| KOY8039 | BY4741 *TEF<sup>P</sup>-mito-DHFR-mCherry::CgHIS3* *get1::natNT2* *FAR8-3×GFP::hphNT1*<br>*far10::kanMX6* |  |
| KOY8045 | BY4741 *TEF<sup>P</sup>-mito-DHFR-mCherry::CgHIS3* *atg32::KIURA3* *atg32(Δ151-200)-3HA* |  |
| KOY8046 | BY4741 *TEF<sup>P</sup>-mito-DHFR-mCherry::CgHIS3* *get1::natNT2*<br>*atg32::KIURA3* *ATG32-3HA* |  |
| KOY8047 | BY4741 *TEF<sup>P</sup>-mito-DHFR-mCherry::CgHIS3* *get1::natNT2*<br>*atg32::KIURA3* *atg32(Δ151-200)-3HA* |  |
| KOY8049 | BY4741 *TEF<sup>P</sup>-mito-DHFR-mCherry::CgHIS3* *get2::natNT2*<br>*atg32::KIURA3* *ATG32-3HA* |  |
| KOY8050 | BY4741 *TEF<sup>P</sup>-mito-DHFR-mCherry::CgHIS3* *get2::natNT2*<br>*atg32::KIURA3* *atg32(Δ151-200)-3HA* |  |
| KOY8051 | BY4741 *TEF<sup>P</sup>-mito-DHFR-mCherry::CgHIS3* *FAR8-3×GFP::natNT2*<br>*far9::KIURA3* |  |
| KOY8054 | BY4741 *TEF<sup>P</sup>-mito-DHFR-mCherry::CgHIS3* *get1::natNT2* *FAR8-3×GFP::hphNT1*<br>*far9::KIURA3* |  |
| KOY8066 | BY4741 *TEF<sup>P</sup>-mito-DHFR-mCherry::CgHIS3* *far10::KIURA3* |  |
| KOY8069 | BY4741 *TEF<sup>P</sup>-mito-DHFR-mCherry::CgHIS3* *get1::natNT2* *far10::natNT2* |  |
| KOY8072 | BY4741 *TEF<sup>P</sup>-mito-DHFR-mCherry::CgHIS3* *get2::natNT2* *far10::natNT2* |  |
| KOY8075 | BY4741 *TEF<sup>P</sup>-mito-DHFR-mCherry::CgHIS3* *atg32::KIURA3* *ATG11-HA-SmBiT::hphNT1*<br>*ATG32-3HA-3×mGFP-3FLAG-LgBiTn* *ppg1::KIURA3* *PPG1* |  |
| KOY8076 | BY4741 *TEF<sup>P</sup>-mito-DHFR-mCherry::CgHIS3* *atg32::KIURA3* *ATG11-HA-SmBiT::hphNT1*<br>*ATG32-3HA-3×mGFP-3FLAG-LgBiTn* *ppg1::KIURA3* *PPG1(H111N)* |  |
| KOY8077 | BY4741 *TEF<sup>P</sup>-mito-DHFR-mCherry::CgHIS3* *FAR8-3×GFP::kanMX6* |  |
| KOY8079 | BY4741 *TEF<sup>P</sup>-mito-DHFR-mCherry::CgHIS3* *get1::natNT2* *FAR8-3×GFP::kanMX6* |  |
| KOY8081 | BY4741 *TEF<sup>P</sup>-mito-DHFR-mCherry::CgHIS3* *get2::natNT2* *FAR8-* |  |
|  | *3×GFP::kanMX6* |  |
| KOY8092 | BY4741 *TEF<sup>P</sup>-mito-DHFR-mCherry::CgHIS3 atg32::KIURA3 ATG11-HA-SmBiT::hphNT1 ATG32-3HA-3×mGFP-3FLAG-LgBiTn ppg1::KIURA3 PPG1 get1::natNT2* |  |
| KOY8095 | BY4741 *TEF<sup>P</sup>-mito-DHFR-mCherry::CgHIS3 atg32::KIURA3 ATG11-HA-SmBiT::hphNT1 ATG32-3HA-3×mGFP-3FLAG-LgBiTn ppg1::KIURA3 PPG1 get2::natNT2* |  |
| KOY8098 | BY4741 *TEF<sup>P</sup>-mito-DHFR-mCherry::CgHIS3 atg32::KIURA3 ATG11-HA-SmBiT::hphNT1 ATG32-3HA-3×mGFP-3FLAG-LgBiTn ppg1::KIURA3 PPG1(H111N) get1::natNT2* |  |
| KOY8101 | BY4741 *TEF<sup>P</sup>-mito-DHFR-mCherry::CgHIS3 atg32::KIURA3 ATG11-HA-SmBiT::hphNT1 ATG32-3HA-3×mGFP-3FLAG-LgBiTn ppg1::KIURA3 PPG1(H111N) get2::natNT2* |  |
| KOY8107 | BY4741 *TEF<sup>P</sup>-mito-DHFR-mCherry::CgHIS3 atg32::KIURA3 ATG11-HA-SmBiT::hphNT1 ATG32-3HA-3×mGFP-3FLAG-LgBiTn far10::KIURA3* |  |
| KOY8110 | BY4741 *TEF<sup>P</sup>-mito-DHFR-mCherry::CgHIS3 atg32::KIURA3 ATG11-HA-SmBiT::hphNT1 ATG32-3HA-3×mGFP-3FLAG-LgBiTn get1::natNT2 far10::KIURA3* |  |
| KOY8113 | BY4741 *TEF<sup>P</sup>-mito-DHFR-mCherry::CgHIS3 atg32::KIURA3 ATG11-HA-SmBiT::hphNT1 ATG32-3HA-3×mGFP-3FLAG-LgBiTn get2::natNT2 far10::KIURA3* |  |
| KOY8115 | BY4741 *TEF<sup>P</sup>-mito-DHFR-mCherry::CgHIS3 atg32::KIURA3 ATG11-HA-SmBiT::hphNT1 ATG32-3HA-3×mGFP-3FLAG-LgBiTn far9::KIURA3 get2::natNT2* |  |
| KOY8179 | BY4741 *TEF<sup>P</sup>-mito-DHFR-mCherry::CgHIS3 FAR8-3×GFP::hphNT1 FAR9-TM<sup>ER</sup>::kanMX6* |  |
| KOY8181 | BY4741 *TEF<sup>P</sup>-mito-DHFR-mCherry::CgHIS3 get1::natNT2 FAR8-3×GFP::hphNT1 FAR9-TM<sup>ER</sup>::kanMX6* |  |
| KOY8183 | BY4741 *TEF<sup>P</sup>-mito-DHFR-mCherry::CgHIS3 get2::natNT2 FAR8-3×GFP::hphNT1 FAR9-TM<sup>ER</sup>::kanMX6* |  |
| KOY8192 | BY4741 *TEF<sup>P</sup>-mito-DHFR-mCherry::CgHIS3 msp1::KIURA3 MSP1(E193Q)* |  |
| KOY8235 | BY4741 *TEF<sup>P</sup>-mito-DHFR-mCherry::CgHIS3 get3::natNT2 FAR8-3×GFP::kanMX6* |  |
| KOY8347 | BY4741 *TEF<sup>P</sup>-mito-DHFR-mCherry::CgHIS3 atg32::KIURA3 ATG32-3HA msp1::natNT2* |  |
| KOY8350 | BY4741 *TEF<sup>P</sup>-mito-DHFR-mCherry::CgHIS3 atg32::KIURA3 atg32(Δ151-200)-3HA msp1::natNT2* |  |
| KOY8353 | BY4741 *TEF<sup>P</sup>-mito-DHFR-mCherry::CgHIS3 atg32::KIURA3 ATG32-3HA get3::natNT2* |  |
| KOY8356 | BY4741 *TEF<sup>P</sup>-mito-DHFR-mCherry::CgHIS3 atg32::KIURA3 atg32(Δ151-200)-3HA get3::natNT2* |  |
| KOY8386 | BY4741 *TEF<sup>P</sup>-mito-DHFR-mCherry::CgHIS3 msp1::natNT2 ppg1::zeoNT3* |  |
| KOY8389 | BY4741 *TEF<sup>P</sup>-mito-DHFR-mCherry::CgHIS3 get3::natNT2 ppg1::zeoNT3* |  |
| KOY8392 | BY4741 *TEF<sup>P</sup>-mito-DHFR-mCherry::CgHIS3 get3::natNT2 msp1::KIURA3 ppg1::zeoNT3* |  |
| KOY8401 | BY4741 *TEF<sup>P</sup>-mito-DHFR-mCherry::CgHIS3 atg32::KIURA3 ATG32-3HA msp1::natNT2 get3::zeoNT3* |  |
| KOY8404 | BY4741 *TEF<sup>P</sup>-mito-DHFR-mCherry::CgHIS3 atg32::KIURA3 atg32(Δ151-200)-3HA msp1::natNT2 get3::zeoNT3* |  |
| KOY8440 | BY4741 *TEF<sup>P</sup>-mito-DHFR-mCherry::CgHIS3 atg32::KIURA3 ATG11-HA-SmBiT::hphNT1 ATG32-3HA-3×mGFP-3FLAG-LgBiTn FAR9-TM<sup>ER</sup>::kanMX6* |  |
| KOY8442 | BY4741 *TEF<sup>P</sup>-mito-DHFR-mCherry::CgHIS3 atg32::KIURA3 ATG11-HA-SmBiT::hphNT1 ATG32-3HA-3×mGFP-3FLAG-LgBiTn get1::natNT2 FAR9-TM<sup>ER</sup>::kanMX6* |  |
| KOY8444 | BY4741 *TEF<sup>P</sup>-mito-DHFR-mCherry::CgHIS3 atg32::KIURA3 ATG11-HA-SmBiT::hphNT1 ATG32-3HA-3×mGFP-3FLAG-LgBiTn get2::natNT2 FAR9-TM<sup>ER</sup>::kanMX6* |  |
| KOY8446 | BY4741 *TEF<sup>P</sup>-mito-DHFR-mCherry::CgHIS3 msp1::natNT2 FAR8-3×GFP::kanMX6* |  |
| KOY8554 | BY4741 *TEF<sup>P</sup>-mito-DHFR-mCherry::CgHIS3 FAR8-3×GFP::kanMX6 atg39::KIURA3 atg40::zeoNT3* |  |
| KOY8557 | BY4741 *TEF<sup>P</sup>-mito-DHFR-mCherry::CgHIS3 FAR8-3×GFP::hphNT1 atg32::natNT2 atg39::KIURA3 atg40::zeoNT3* |  |
| KOY8568 | BY4741 *TEF<sup>P</sup>-mito-DHFR-mCherry::CgHIS3 FAR8-3×GFP::natNT2 FAR9-TA<sup>MITO</sup>::hphNT1 FAR10-TA<sup>MITO</sup>::kanMX6* |  |
| KOY8570 | BY4741 *TEF<sup>P</sup>-mito-DHFR-mCherry::CgHIS3 ppg1::KIURA3 PPG1 FAR8-3×GFP::natNT2 FAR9-TA<sup>MITO</sup>::hphNT1 FAR10-TA<sup>MITO</sup>::kanMX6* |  |
| KOY8572 | BY4741 *TEF<sup>P</sup>-mito-DHFR-mCherry::CgHIS3 ppg1::KIURA3 PPG1(H111N) FAR8-3×GFP::natNT2 FAR9-TA<sup>MITO</sup>::hphNT1 FAR10-TA<sup>MITO</sup>::kanMX6* |  |
| KOY8719 | BY4741 *TEF<sup>P</sup>-mito-DHFR-mCherry::CgHIS3 msp1::KIURA3 MSP1(E193Q) get3::natNT2* |  |
| KOY8793 | BY4741 *SEC63-mCherry::KIURA3 FAR8-3×GFP::hphNT1 FAR9-TM<sup>ER</sup>::kanMX6* |  |
| KOY8795 | BY4741 *SEC63-mCherry::KIURA3 get1::natNT2 FAR8-3×GFP::hphNT1 FAR9-TM<sup>ER</sup>::kanMX6* |  |
| KOY9095 | BY4741 *SEC63-mCherry::KIURA3 get2::natNT2 FAR8-3×GFP::kanMX6 FAR9-TM<sup>ER</sup>::hphNT1* |  |

1. Brachmann, C.B., Davies, A., Cost, G.J., Caputo, E., Li, J., Hieter, P. and Boeke, J.D. (1998). Designer deletion strains derived from *Saccharomyces cerevisiae* S288C: a useful set of strains and plasmids for PCR-mediated gene disruption and other applications. *Yeast* **14**, 115-132.

**Table S2.** Plasmid used in this study.

| Plasmid number | Name | Relevant characteristics |
| --- | --- | --- |
| KOB71 | pRS316-ATG32-3HAn | *CEN URA3 580 bp 5'-UTR & 744 bp 3'-UTR from ATG32* |

## Notes

### Competing Interest Statement

The authors have declared no competing interest.

